# Integrative genome-wide analysis reveals EIF3A as a key downstream regulator of translational repressor protein Musashi 2 (MSI2)

**DOI:** 10.1101/2021.02.06.428911

**Authors:** Shilpita Karmakar, Oscar Ramirez, Kiran V. Paul, Abhishek K. Gupta, Valentina Botti, Igor Ruiz de los Mozos, Nils Neuenkirchen, Robert J. Ross, Karla M. Neugebauer, Manoj M. Pillai

## Abstract

Musashi 2 (MSI2) is an RNA binding protein (RBP) that regulates asymmetric cell division and cell fate decisions in normal and cancer stem cells. MSI2 appears to repress translation by binding to 3’ untranslated regions (3’UTRs) of mRNA, but the identity of functional targets remains unknown. Here we used iCLIP to identify direct RNA binding partners of MSI2 and integrated these data with polysome profiling to obtain insights into MSI2 function. iCLIP revealed specific MSI2 binding to thousands of target mRNAs largely in 3’UTRs, but translational differences were restricted to a small fraction of these transcripts, indicating that MSI2 regulation is not triggered by simple binding. Instead, the functional targets identified here were bound at higher density and contain more “U/TAG” motifs compared to targets bound non-productively. To further distinguish direct and indirect targets, MSI2 was acutely depleted. Surprisingly, only 50 transcripts were found to undergo translational induction on acute MSI2 loss. Eukaryotic elongation factor 3A (EIF3A) was determined to be an immediate, direct target. We propose that MSI2 down-regulation of EIF3A amplifies these effects on the proteome. Our results also underscore the challenges in defining functional targets of RBP since mere binding does not imply a discernible functional interaction.

## INTRODUCTION

RNA binding proteins (RBP) encompass a diverse group of proteins that regulate all aspects of RNA biology. The Musashi proteins (MSI1 and its homologue MSI2) are highly conserved across metazoans and contain two distinct RNA recognition motifs (RRM) (1). MSI proteins are thought to bind to the 3’ untranslated region (3’UTR) of specific transcripts and regulate their translation. Of the two Musashi homologues, MSI1 is expressed primarily in neurons (2). In contrast, MSI2 is ubiquitous in its expression. MSI2 is expressed at high levels in hematopoietic stem and progenitor cells, where its down-regulation coincides with stem cell differentiation (3). A role for deregulated MSI2 expression in cancer was first reported in aggressive myeloid leukemia. MSI2 was shown to be transcriptionally up-regulated in both chronic myelogenous leukemia in myeloid blast crisis (CML-BC) and aggressive acute myeloid leukemia (AML) (3,4). High expression of both MSI homologues have since been reported in other malignancies such as colorectal cancer (5), pancreatic cancer (6), medulloblastoma (7), breast cancer (8) and lung cancer (9).

The critical role of MSI proteins in regulating asymmetric cell division and cell fate suggests a mechanism that regulates a small number of specific molecular targets. In early studies without genome-wide analyses, it was proposed that NUMB, an inhibitor of Notch signaling, is a critical target (3,10). However, later studies were not been able to confirm this association in neural, blood or lung cancer cells (9–12). A number of genome-wide studies have defined the RNA-interactome of MSI1 and MSI2 in multiple cell types. These include CLIP-Seq (or HITS-CLIP) in mouse keratinocytes, leukemic cell lines, embryonic kidney cell lines and intestinal epithelium (11,13–15). Complementary techniques such as SELEX (systematic evolution of ligands by exponential enrichment) and TRIBE (targets of RNA-binding proteins identified by editing) have also been used to determine targets of MSI proteins (16,17). These studies demonstrated that MSI proteins bind to thousands of transcripts in a cell context-specific manner. While many functionally relevant targets were noted to be bound by MSI homologues in each of these studies, unbiased analysis of functional regulation of these bound targets was not performed. Hence these studies are limited in providing in-depth understanding of how MSI proteins regulate gene expression.

In this study, we sought to answer this question: Which of these bound targets are translationally modulated by MSI proteins? We hypothesized that MSI2 affects the translation of only a subset of the transcripts it binds to. To test this, we integrated two genome-wide approaches-individual nucleotide resolution Cross-Linking and Immunoprecipitation (iCLIP) and polysome profiling to address the relationship between MSI2 binding and translational regulation. Because we are interested in understanding the role of MSI2 in cancers where MSI2 is expressed, we established FLAG-tagged MSI2 expressed in K562 cells as a model system. The K562 cell line was derived from the blast crisis stage of CML patient and has high constitutive expression of MSI2 (11,12). Analysis of the data reveals that although MSI2 binds to the 3’UTR of over four thousand transcripts in this study, only a fraction (2.6%) of these are translationally regulated. Through acute depletion of MSI2 and polysome profiling, we also identify EIF3A as a critical downstream regulator of MSI2. Additionally, our results argue for the need to incorporate functional assays intandem with CLIP-Seq approaches to distinguish binding and regulatory functions of an RBP.

## MATERIALS AND METHODS

### Cell culture

Cell lines were obtained from ATCC. K562 cells (12), and derivatives were cultured in RPMI-1640 supplemented with 10% fetal bovine serum (FBS). HEK293T and NIH3T3 cells were grown in DMEM supplemented with 10% FBS. Puromycin selection (for stable FLAG-MSI2 overexpression or shRNA mediated knockdown) was performed at 1 μg/ml concentration. Neomycin (G418) selection (for inducible shRNA clones) was performed at 800μg/ml concentration. Single cell clones were selected after 10-14 days of selection by plating cells in methyl-cellulose as previously described (18)

### Cloning, plasmid constructs and viral vector production

The human MSI2 ORF (NM_13892.2) was PCR amplified from cDNA and cloned into the BamHI and EcoRI sites of the pBABE-puro retroviral vector with an N-terminal FLAG tag. Stable lentiviral vectors expressing shRNA targeting MSI2 and control shRNAs were obtained from Sigma Aldrich (Mission lentiviral system, based on the pLKO.1 vector; Clone details are provided in **Supplementary Methods**) and confirmed for their knockdown activity by RT-PCR and Western blotting. Lentiviral vector pLKO-Tet-On (Addgene Plasmid #21916) was used to generate inducible knockdown clones of MSI2 and EIF3A (19) (**for the sequence see Supplementary Methods**). Retroviral and lentiviral vectors were produced by the co-transfection of respective pro-viral plasmids with appropriate helper and envelope plasmids and transduced into K562 cells as previously described (18).

### iCLIP for MSI2

iCLIP for FLAG-MSI2 was performed as previously reported (20) with minor variations (21). Briefly, 40 million K562 cells expressing FLAG-MSI2 were cross-linked twice (4 mJ and 2 mJ pulses) and stored at −80°C prior to analysis. Three individual clones (three single cell clones of FLAG-MSI2). RNA crosslinking to FLAG-MSI2 was confirmed and RNAse conditions were optimized prior to library preparation (**Supplementary Figures 1A-D**). mRNA-Seq libraries were prepared from RNA isolated from corresponding batches of cells. For both iCLIP and mRNA-Seq, 50 base-pair, single-end sequencing was performed on the Illumina HiSeq2000 platform.

### Polysome profiling

Polysome profiling was performed using single cell clones of K562 as previously reported (22,23) and details are in **Supplementary Methods**. Briefly, ~ 40 million cells in log-phase growth were treated with cycloheximide (1 μg/ml) for 10 minutes, lysed in TMK-lysis buffer, cleared of debris by centrifugation and loaded on a 10-60% sucrose gradient. Polysome fractionation was achieved by ultracentrifugation and individual fractions were collected (46 fractions of approximately 800 μl) using the Teldyne ISCO automated fraction collector with continuous monitoring of the absorbance at 254 nm. Fractions corresponding to heavier polysomes were pooled. Total and polysomal RNAs were isolated by Trizol (Life Technologies, Carlsbad, CA). RNA-Seq libraries for polysomal RNA and total RNA were prepared using the Illumina TruSeq kit and were sequenced on the Illumina HiSeq2000 (single-end, 50 base-pair). A detailed protocol for polysome profiling is provided in the **Supplementary Methods**.

### Luciferase assay

The 3’UTR of EIF3A (or mutants lacking U/TAG motif) were cloned downstream of the Renilla luciferase in the XhoI and NotI sites of psiCHECK2 vector (Promega, Madison, WI; sequence details provided in the **Supplementary Methods**). 40ng of the psiCHECK2 plasmid was transiently co-transfected with 320ng of MSI2-pCDNA3.1(+) in NIH3T3 (24) cells using ITX2 transfection reagent and luciferase activity (renilla and firefly) was measured 24hours later with the Dual Glow-Stop and Glow luciferase kit (Promega) using a BioTek luminometer (Synergy).

### Western blot analysis

For Western blot analysis, cells were lysed using 1X RIPA buffer (10 mM Tris-HCl (pH 8.0), 1 mM EDTA, 0.5 mM EGTA, 1% Triton X-100, 0.1% sodium deoxycholate) supplemented with 1X complete mini EDTA protease inhibitor cocktail (Roche), incubated on ice for 15 minutes, and the supernatant was isolated by centrifugation. Protein concentration was determined using the DC protein assay (Bio-Rad) following the manufacturer’s recommendations. 30μg of total protein was resolved on 10% precast SDS-PAGE gel (Bio-Rad) and transferred to a methanol-preconditioned PVDF membrane by wet transfer for 90 minutes at 100V. Membranes were blocked with 5% nonfat dry milk and probed with the appropriate primary and secondary antibodies. Immunoreactive bands were visualized using Electro Chemiluminescence (ECL, Roche). Details of antibodies used and other conditions for blotting are summarized in **Supplementary Methods**.

### Bioinformatic analysis and Statistics

Details of the bioinformatic analysis are provided in **Supplementary Methods**. The Mann-Whitney non-parametric testing was used to determine statistical significance for comparisons of next generation sequencing datasets. Overlap between datasets was performed using Fisher’s exact test (Gene overlap function of R package Bioconductor). Students t-test was used for other comparisons.

## RESULTS

### The RNA interactome of MSI2 in K562 cells

Like other protocols that detect RNA-protein interactions, iCLIP utilizes ultraviolet (UV) radiation to cross-link RNA to adjacent protein moieties at 0 Å, allowing stringent immunoprecipitation (IP) of RNA-protein complexes (25). Three single-cell clones of K562 cells expressing FLAG-tagged MSI2 were isolated and verified for stable expression of FLAG-MSI2 (**Supplementary Figure 1A**). FLAG-MSI2 cross-linking to RNA by UV was confirmed (**Supplementary Figure 1B**) and RNAse A digestion was optimized to attain the optimal distribution of the MSI2-RNA smear (above the predicted molecular weight of FLAG-MSI2 (37kD) (**Supplementary Figure 1C**). Illumina-compatible libraries were prepared from the isolated RNA (**Materials and Methods & Supplementary Figure 1D**) and sequenced to a depth of about 50 million single end reads per sample. PCR duplicates were eliminated by introducing five base-pair random sequences during the reverse transcription step as unique molecular identifiers (UMIs). After mapping to the human genome (hg19), cross-link sites and clusters were determined by the iCount algorithm (26), http://icount.biolab.si/. In iCLIP, cDNA start sites are annotated as cross-link sites, and regions with significant clustering of cross-link sites are designated as “cross-link clusters” (25). Crosslink clusters that met our statistical cut-off (FDR < 0.05) and were represented in at least two of three biological replicates were designated as high confidence clusters and used for further analysis. A high degree of overlap was found between the target genes identified between the three biological replicates (**Figure 1A**). The positive correlation (R^2^ =0.625) between transcript abundance and iCLIP abundance (**Figure 1B**) is typical of iCLIP experiments, given that RNAs must be expressed to be detected; the distribution confirms that a broad range of expression levels are represented in the iCLIP dataset.

**Figure 1.**
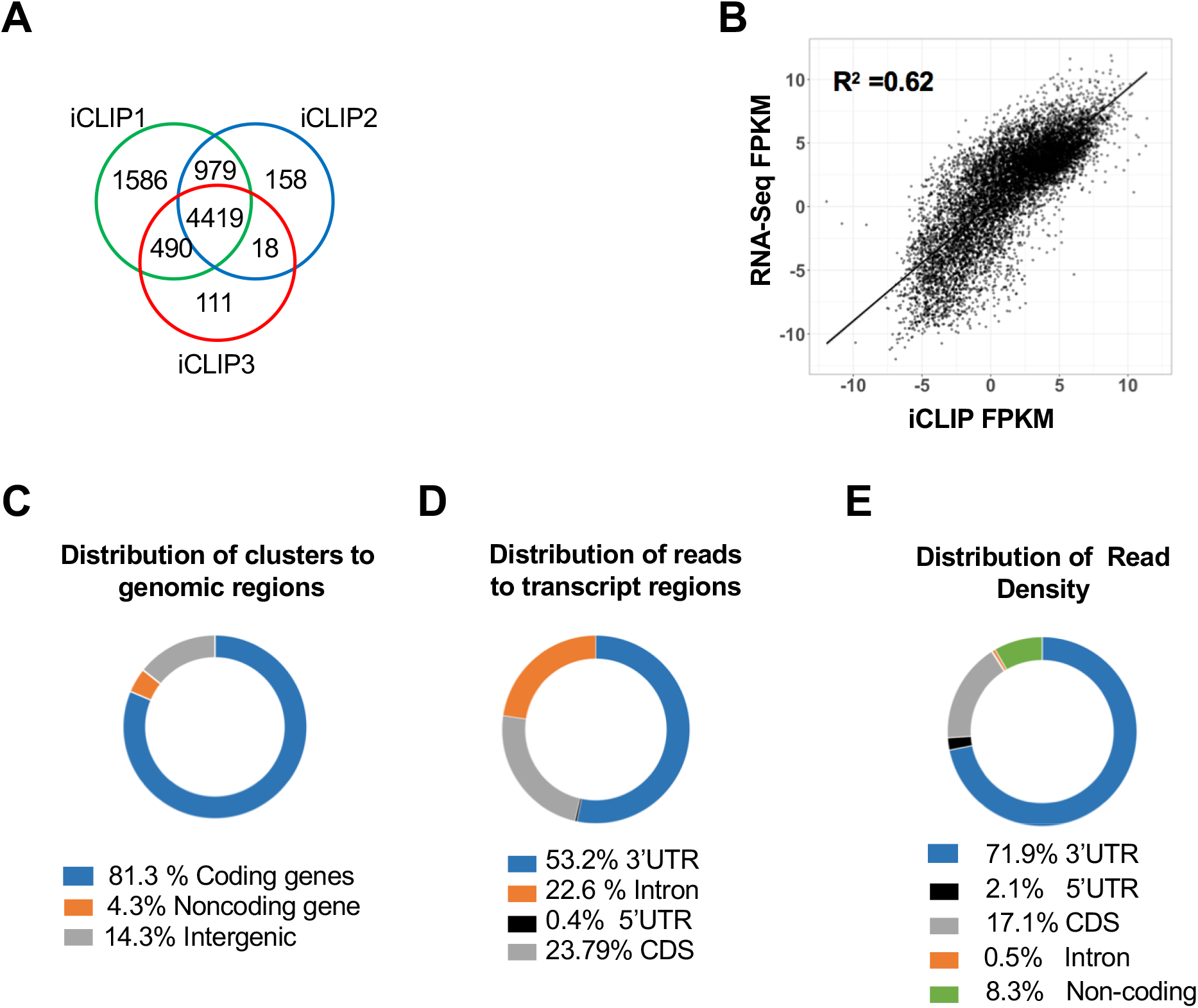
iCLIP analysis of FLAG-MSI2 across genomic and transcriptomic regions. (A) Overlap of transcripts with 3’UTR peaks between three replicate iCLIP experiments. Transcripts with high confidence cross-link clusters determined by iCount were used for analysis. 4419 transcripts were found to have crosslink clusters in 3’UTRs in all three replicates. (B) Scatter plot showing correlation between MSI2 binding (normalized FPKM from three iCLIP experiments) and transcript abundance (normalized RNA-Seq FPKM), R^2^ =0.6245. (C) Distribution of aligned iCLIP clusters across genomic regions (protein-coding genes, noncoding genes, and intergenic regions). Pie chart plotted as a percentage for each region (total 100%). (D) Distribution of iCLIP cross-link clusters across different transcript regions (5’UTR, CDS, 3’UTR and introns). Pie chart plotted as a percentage for each region (total 100%). (E) Distribution of absolute read counts normalized to the length of each transcript region. Total number of reads that were uniquely aligned to each of the regions was determined (FPKM). This was normalized to the total length of those regions. Pie chart plotted as a percentage for each region (total 100%).

Most cross-link clusters (81.3%) were identified in protein-coding genes, with noncoding-genes and intergenic regions encompassing only 4.3% and 14.3% of clusters respectively (**Figure 1C**). Within protein-coding genes, clusters were enriched in 3’UTR (53.2%), in agreement with the purported role of MSI2 as a 3’UTR binding protein (**Figure 1D**). Fewer than 1% of clusters were localized to 5’UTR, and 23.8% were in protein coding regions (CDS). 22.6% of clusters were noted to be localized to introns. The density of these clusters (normalized to the respective length of the transcript region) is shown in **Figure 1E**. Such normalized density was by far the highest in 3’UTR (71.9%) followed by CDS (17.1%). Introns had the lowest density (0.5%) despite having highest total reads aligning to it. The large proportion of total iCLIP reads aligning to introns was unexpected, given that MSI proteins are thought to be cytoplasmic in location due to their primary role in translational repression. However, recent reports have suggested a nuclear localization for MSI2 during specific phases of the cell cycle (27) and associated with neurodegenerative disorders (28). Reassociation of RNA binding proteins with target RNA after cell lysis has been reported (29) which may represent another source of intronic reads. Intronic reads have also been reported for MSI1 CLIP-Seq (30). Taken together, our results suggest predominant binding of MSI2 to the 3’UTR region of protein-coding transcripts, with sparse binding to other regions including introns.

### MSI2 binding motifs are enriched for the U/TAG motif and poly-U/T motifs

Previous studies using genome-wide profiling of MSI2-bound RNA or SELEX (Systematic evolution of ligands by exponential enrichment) have shown that MSI proteins bind to UAG in RNA and TAG in DNA (16,31, 32) or contains poly-U/T stretches (14,30). We implemented two strategies to search for motif enrichment in the iCLIP datasets. We first examined the enrichment of motifs in a window from −20 to +20 base pairs from the high confidence cross-link sites using the HOMER algorithm (33). Motifs thus enriched typically included “U/TAG” containing motifs across the different transcriptomic regions (**Figure 2A**). We then examined enrichment of specific pentamers in the cross-link clusters, as determined by iCount (25). Notably, these pentamers revealed enrichment for U/TAG motifs or high U/T content (**Figure 2B**). Finally, we determined the distribution of the U/TAG motif and U/T-rich sequences from high confidence cross-link sites (**Figures 2C and 2D** respectively). These motif features were found enriched around the crosslink sites. Combined, our results confirm an enrichment of U/TAG or polyU/T containing motifs in MSI2 binding sites. A modest bias towards uracil-containing stretches is characteristic of CLIP dataset due to preferential crosslinking (34). However, AG should not reflect a bias in the crosslinking, so U/TAG-containing sequences do appear to represent the preferred binding motifs for MSI2.

**Figure 2.**
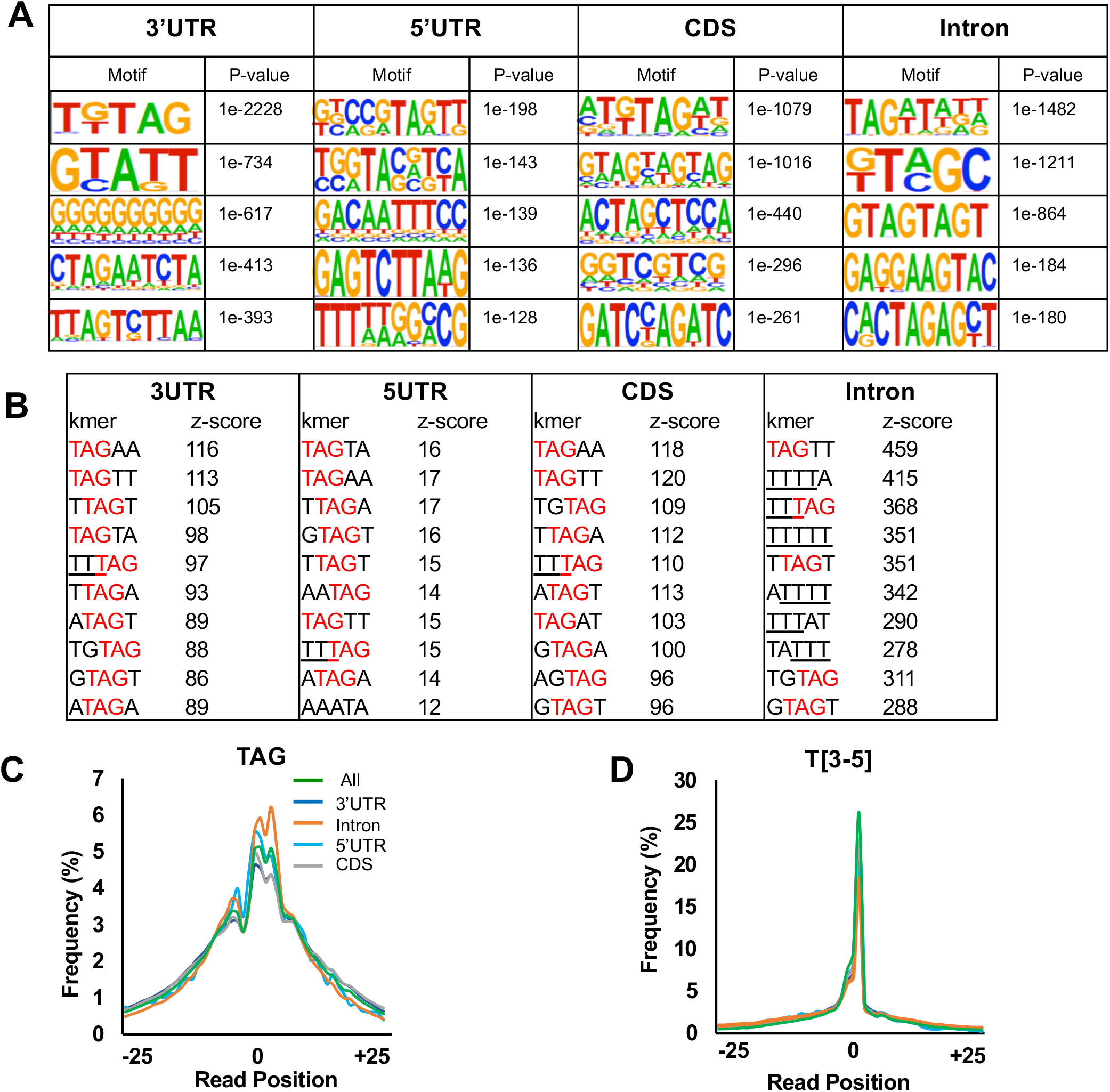
Distribution of FLAG-MSI2 iCLIP start sites and enrichment of TAG and poly-T containing 5pentamers and motifs. (A) Motif identified in regions −10 to +10 from high confidence cross-link sites (FDR<0.05) by HOMER algorithm. Top 5 motifs along with the corresponding p-value are shown for each of the transcript regions (5’UTR, CDS, 3’UTR and introns). (B) Ten most frequent 5-mers identified in a window of −10 to +10 nucleotides with respect to cross-link sites for each of the transcript regions (5’UTR, CDS, 3’UTR and introns). TAG within these 5-mers are denoted in red and poly-T stretches (more than 3) are underlined. (C) Distribution of “TAG” motif upstream and downstream (−25 to +25 nucleotides) of the crosslink sites. Frequency distribution across each transcriptomic region (5’UTR, CDS, 3’UTR and introns) along with aggregate distribution across all regions (“All”) is shown. Enrichment of TAG motif is seen around cross-link site. (D) Distribution of poly-T motifs (3 or more) with respect to cross-link site positions, plotted in a similar fashion as Figure 2C. Enrichment for poly-T stretches are seen around the cross-link sites.

### Polysome profiling reveals distinct effects on translatome compared with transcriptome

Specific binding of MSI2 to thousands of targets as revealed by iCLIP was surprising, given the specific biological roles of MSI2 on cell-fate decisions. We hypothesized that only a subset of target transcripts bound by MSI2 actually undergo functionally relevant translational regulation, and thus be defined as translational targets of “productive” MSI2 binding. To define this subset, we first generated stable MSI2 knockdown (MSI2-KD) and control cells with lentiviral shRNA constructs. After verifying reduction of MSI2 (**Figure 3A and Supplementary Figure 2**), polysome profiling was performed (22). Polyribosomes or polysomes are aggregates of two or more ribosomes assembled on mRNA undergoing efficient translation (35). By comparing the change in abundance of transcripts associated with polysomes to the change in total transcript levels, changes in translation can be inferred (36). To perform polysome profiling, we generated single cell clones of MSI2 knockdown K562 cells (MSI2-KD) with stable lentiviral expression of shRNA, as well as control clones with scrambled shRNA (**Figure 3A**). Polysome and transcriptome profiles were generated for each of these clones in three replicates, (**Figures 3B-C**). Potential productive translational targets of MSI2 were identified as those transcripts that changed at least two-fold by polysome profiling without significant changes in total cellular RNA, as described previously (37). By this criterion, a total of 597 high-confidence genes were identified, with 388 RNA increased in polysomes in response to MSI2 knock-down and 209 decreased (**Figure 3D, 3F & Supplementary File 1**). In contrast, only 103 genes changed by total expression levels (**Figure 3E, 3G and Supplementary File 1**). Concomitantly, transcripts with altered transcript abundance in polysome versus total RNA fractions were poorly correlated (R^2^ = 0.237, **Figure 3H**). In all, 2.6% of genes underwent translational change (defined as altered polysome-specific mRNA abundance) while only 0.46% changed transcriptionally. Together, our results suggest a distinct role for MSI2 in translation regulation.

**Figure 3.**
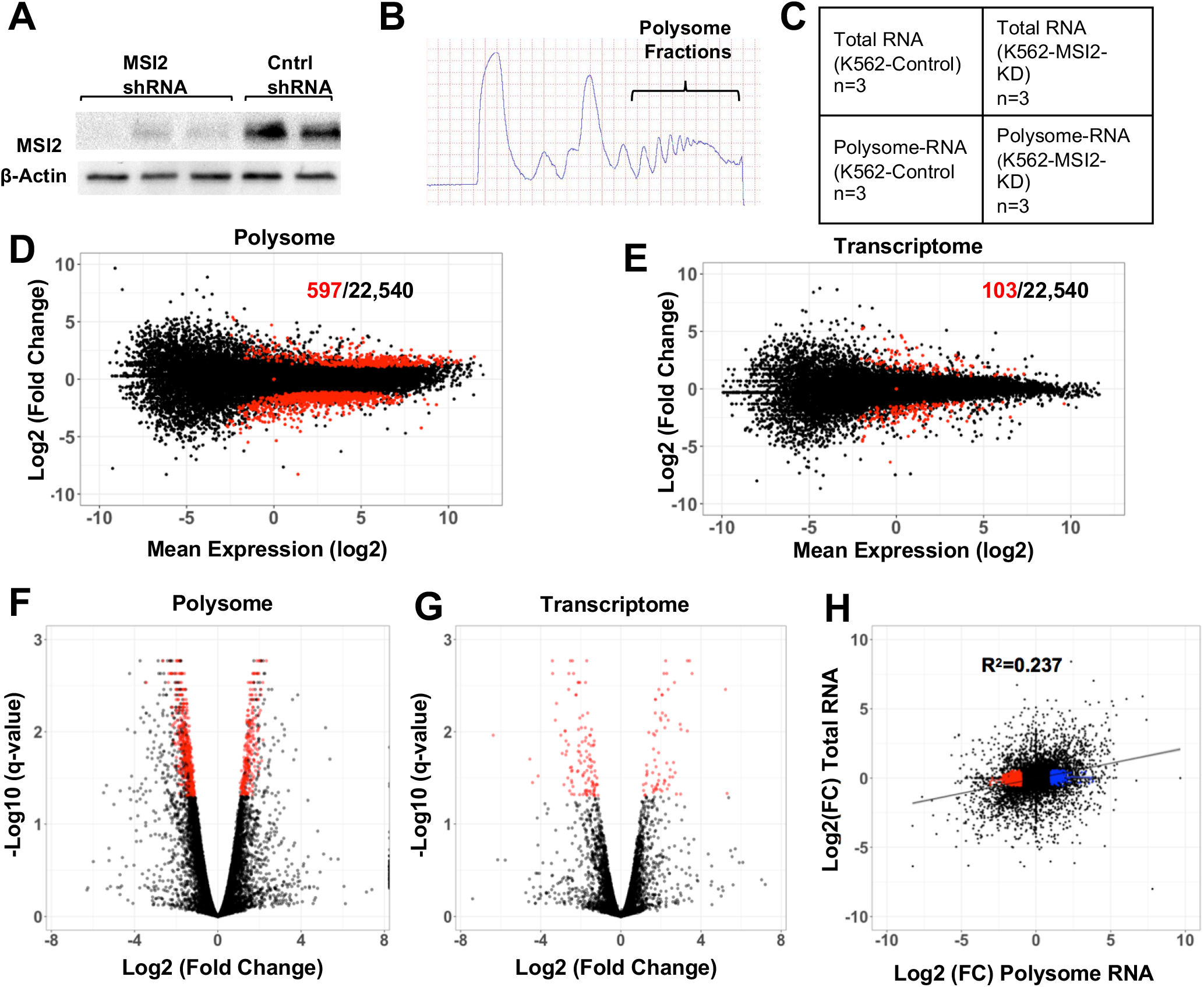
Polysome analysis of MSI2-knockdown (KD) and control cells. (A) Western blot of MSI2-KD and control K562 cells showing loss of MSI2 expression by stable expression of anti-MSI2 shRNA. (B) Absorbance at 254 nm of sucrose gradients for polysome isolation. Higher density fractions as indicated were pooled for RNA isolation. (C) Experimental scheme for polysome analysis. RNA-Seq was performed on paired polysomal RNA and total RNA from wild-type and MSI2-KD cells (in three biological replicates). (D) MA plot (Log_2_ mean expression plotted against Log_2_ mean expression) for polysome profiling (MSI2 knockdown vs control). Red dots represent 597 transcripts that changed significantly among the 22,540 total transcripts (black dots). (E) MA plot for total cellular RNA, plotted similarly to panel (D). Red dots represent 103 transcripts that change significantly among the 22,540 total transcripts (in black). (F) Volcano plot showing overall changes to gene expression in MSI2 knockdown cells compared to control in polysome fraction. Plotted are the log_2_ fold change and −log_10_ (q-value) for genes analyzed for differential expression between control and MSI2-KD. Red dots correspond to high-confidence genes (FDR <0.05). In the plot, q-value refers to FDR (false discovery rate adjusted p-value). (G) Volcano plot showing overall changes to gene expression in MSI2-KD cells compared control in total fraction, as in panel (F). Red dots correspond to high-confidence genes (FDR <0.05). (H) Scatter plot of changes in transcript abundance, polysome vs total RNA (LogFC2). R2 of dispersion calculated to be 0.237. Blue dots represent genes upregulated and red dots represent those downregulated (upon MSI2 knockdown).

### Integration of iCLIP and polysome profiling identifies high-confidence targets of MSI2

Since polysome profiling was performed in cells with sustained MSI2 knockdown, it is not evident which of the transcripts are direct targets of MSI2 and which are indirectly regulated (through other downstream mediators). To determine direct MSI2 targets, we first cross-referenced our list of MSI2-dependent, translationally-regulated target transcripts with mRNAs with high-confidence 3’UTR iCLIP cross-link clusters (**Supplementary File 2**). While 53.8% genes that were translationally up-regulated in response to MSI2-knockdown had iCLIP clusters within their 3’UTR, only 26.5% of those down-regulated by MSI2-knockdown had similar 3’UTR peaks. A similar proportion (24.8%) of transcripts with no change in translation also had peaks in their 3’UTR (**Figure 4A**). We next sought to determine features that distinguish productive MSI2 targets from non-productive binding events. Three groups of MSI-bound transcripts were identified for detailed analysis: (1) those translationally up-regulated in response to MSI2 knockdown with iCLIP peaks in 3’UTR (UP), (2) those translationally down-regulated by MSI2 knockdown with similar iCLIP peaks (DOWN) and (3) mRNAs translationally unchanged with iCLIP peaks (COMPARABLE). We analyzed several attributes of genes within these subsets to determine what distinguishing features might predict productive binding events, including primary sequence motifs, density of cross-link clusters, density of motifs, and secondary structure constraints.

**Figure 4.**
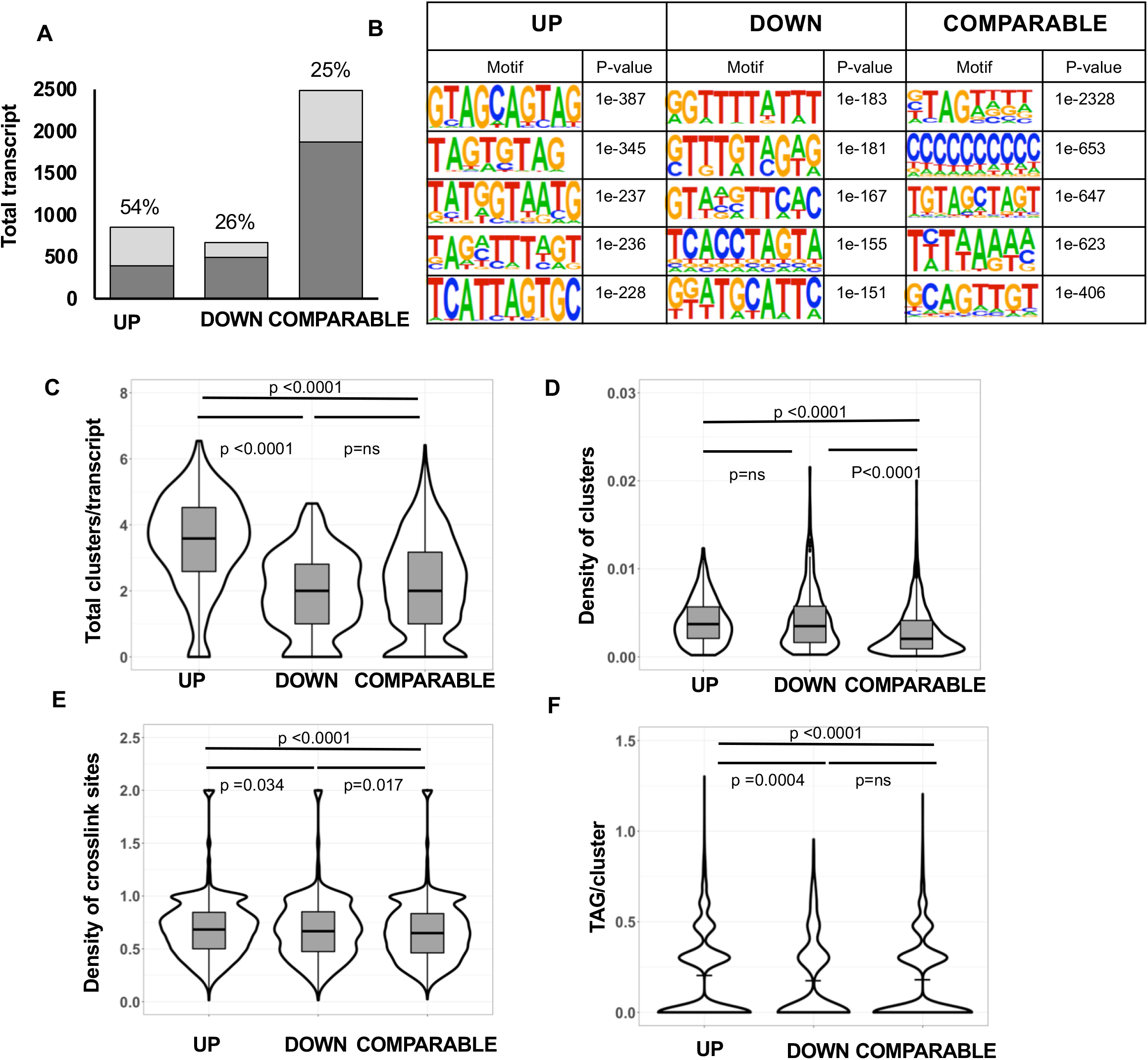
Features of transcripts translationally changing upon MSI2 knockdown and have high confidence iCLIP peaks in 3’UTR. (A) Proportion of genes in the polysome regulated groups that have iCLIP peaks within the transcript regions. 458/851 (54%) of upregulated genes (UP) have iCLIP peaks while only 178/673 (26%) of downregulated genes (DOWN) and 617/2488 (25%) of unchanged genes (COMPARABLE) have similar iCLIP peaks. (B) Motif identified in regions −10 to +10 (with respect to cross-link sites with FDR<0.05) by HOMER algorithm in the upregulated, downregulated or unchanged. Top 5 motifs along with the corresponding P-value are shown for each of the transcript regions (5’UTR, CDS, 3’UTR and introns). (C) Total number of cross-link clusters per transcript among those transcripts with iCLIP clusters. Results segregated in three groups based on polysome profiling results. Box plots within violin plots show mean value as well as p-value determined by Mann-Whitney test. (D) Density of cross-link clusters (total number of high confidence clusters normalized to the length of the transcript) for each of the three groups, plotted similar to that in panel (C). (E) Density of cross-link sites (total number of cross-link sites with FDR <0.05 that falls within the transcript coordinates normalized to length of the transcript) for each of the three groups, plotted similar to panel (C). (F) Total number of TAG motifs (log 2) found within high-confidence crosslink clusters for each of the three groups. The line in the middle of violin plot denotes mean value.

Primary sequence motif analysis in these three subsets showed an enrichment for U/TAG containing sequences in the UP dataset (**Figure 4B**). We next looked at the possibility that productive binding by MSI2 requires multiple molecules binding to the target, which could be inferred from the number and density of cross-link clusters in the iCLIP (25) datasets (cross-link clusters are those regions within iCLIP alignments that cluster together (25). UP targets were found to have a significantly higher number of total cross-link clusters per transcript and density of cross-link clusters normalized to transcript length (**Figures 4C and D** respectively). Additionally, the UP targets also had higher number of individual iCLIP cross-link sites with higher U/TAG motif (**Figure 4E and 4F**). Together, our results show that productive binding of MSI2 binding to down-regulate translation is dependent on high density binding of MSI2. The wide distribution of values in our analysis (**Figures 4C-F**) suggests that productive binding may have additional requirements (such as binding by other RBP).

Given that secondary structure of target transcripts is now known to be a major determinant of RBP-RNA interactions (38), we analyzed the secondary structure of RNA around cross-link sites in the three groups of MSI-bound transcripts using the CapR algorithm (39). We found that crosslink sites or clusters from the three subgroups did not differ from each other with regards to their likelihood to form secondary structures or their relative accessibility (**Supplementary Figures 3A-F**).

### Numerous cancer-relevant genes and pathways are regulated by MSI2

To determine global changes brought about by MSI2 depletion, we performed pathway analysis of transcripts that changed at the level of translation using the ingenuity pathway analysis (IPA) algorithm. Transcripts found translationally up-regulated upon MSI2 knockdown were highly enriched within categories of cancer, cell cycle, cell death and differentiation (**Figure 5A**). Enrichment for pathways in down-regulated genes was less pronounced (lower p values), but somewhat overlapping with the pathways in the up-regulated set (**Figure 5B**). We then analyzed individual transcripts predicted to be up-regulated upon MSI2 knockdown without discernible change in total mRNA levels. These included several encoding cancer-relevant proteins, including EIF3A, MYC, CDK6, SP1, RAD21, USP28, FOXO family proteins and STAT signaling regulators, (full list in **Supplementary File 1**). To determine if protein levels of these transcripts changed as predicted by polysome profiling, we performed Western blot and densitometric analysis by normalizing with β-actin loading control (**Figure 5C, 5D and Supplementary Figure4**). Protein levels generally followed changes predicted from polysome data. Importantly, these transcripts also had putative bindings sites in their respective 3’UTR as shown by iCLIP (**Figure 5C**). Some of the targets (such as C-JUN, CDK1, CTNN1 and RB1) predicted to change per polysome analysis did not have discernible differences (**Supplementary Figure 4**). Our results show that MSI2 affects translation of multiple transcripts related to cancer-related pathways. Given the widespread binding of MSI2 to numerous targets, it is difficult to discern which of these transcripts are directly regulated by MSI2 and not indirect targets.

**Figure 5.**
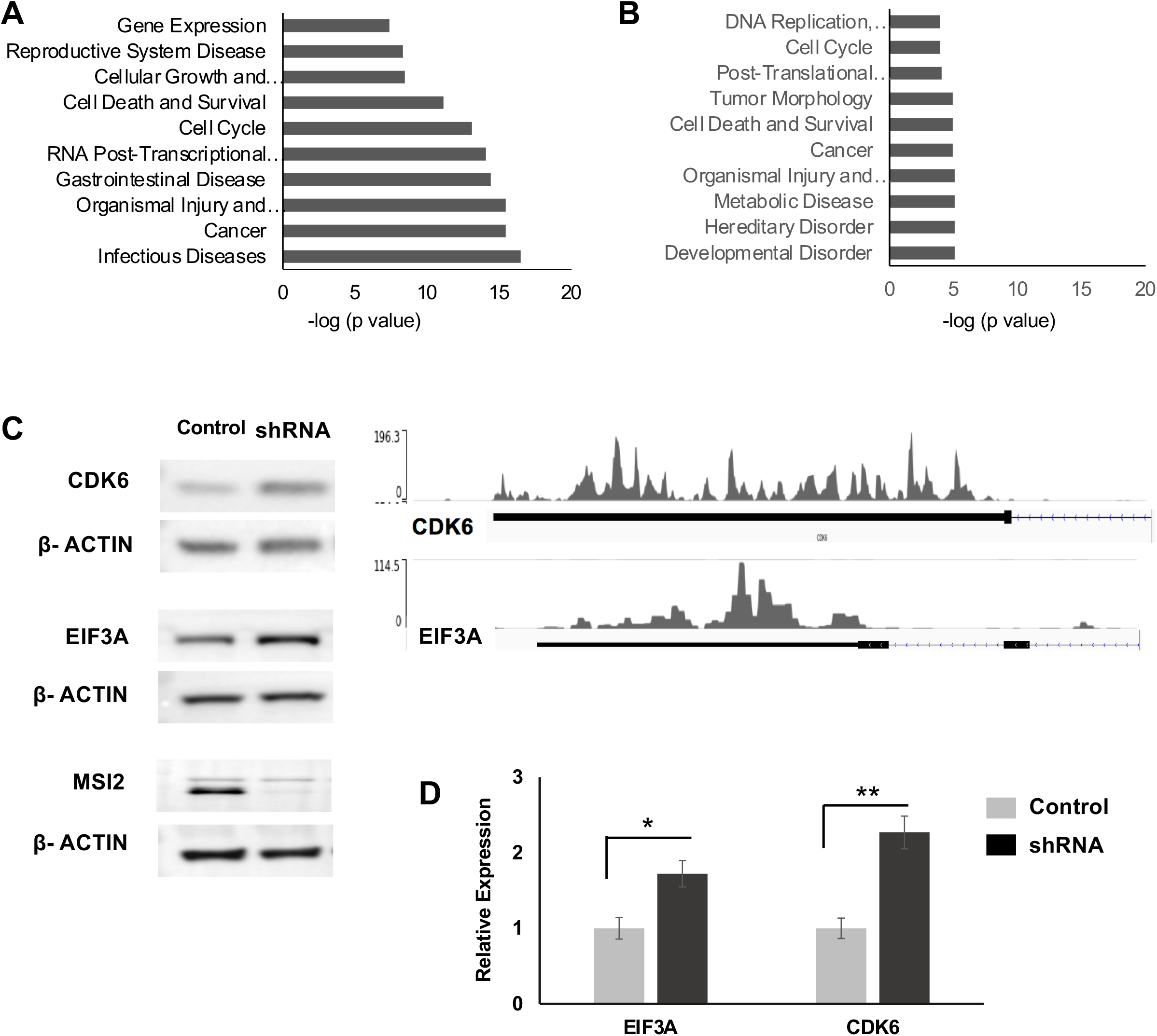
Biological Pathways and genes targeted by MSI2. (A &B) Pathway analysis of MSI2-regulated genes for diseases and functions. The analysis was conducted with high-confidence genes showing differential regulation in polysome fraction (upregulated (A) and downregulated (B)). Plotted are the top 10 pathways and corresponding −log_10_ (p-value). (C) Western blot for candidate genes CDK6 and EIF3A upon stable lentiviral expression of either control (scramble) shRNA, or MSI2 shRNA. Also included are western blots for MSI2 (showing knockdown of MSI2 with shRNA expression) and loading control (β-actin). On the right side of the panel are genome coverage plots of corresponding 3’UTR from iCLIP for MSI2. (D) Quantification of EIF3A and CDK6 Western blots (compared with loading control) from 3 replicates (β-actin). * denotes a p value of 0.0332 and ** a p value of 0.0076

### EIF3A is translationally regulated by MSI2

To further distinguish direct targets of MSI2 from indirect ones, we performed polysome profiling of short-term knockdown of MSI2 in a doxycycline-inducible shRNA system. We speculated that direct targets will have an early effect on translation. We generated single cell clones of tet-inducible shRNA directed against MSI2 with reliable inducible knockdown upon doxycycline addition (> 80% at 48 hours, **Figure 6C and Supplementary Figure 5A, 5B**). Polysome profiling was performed as for the stable MSI2 knockdown described above (comparing induced and uninduced cells) (**Figure 6A and Supplementary Figure 5C,5D**) (22). A total of 50 genes were shown to change significantly at the level of translation. We observed that similar to stable knockdown, the change in polysome was more pronounced than for the transcriptome: only MSI2 itself changed at level of transcription while translation of 50 genes were noted to change (**Figures 6B and Supplementary File 1**). Comparing the datasets (stable vs inducible knockdowns), we noted 10 genes that change in both datasets (**Figure 6B**). Of these, EIF3A was noted to be up-regulated two-fold in the inducible knockdown and 6 fold in stable knockdown suggesting an immediate and sustained effect from MSI2 knockdown. EIF3A (eukaryotic translation initiation factor 3A) is the largest subunit of eIF3, which plays a central role in the recruitment of pre-initiation complex (PIC) to mRNA to initiate peptide translation (40,41). In addition to this role as a canonical regulator of translation, eIF3 components are now recognized to have specialized roles in regulating translation of specific transcripts (42). Other genes in the subset lacked known regulatory functions. Combined, we chose to focus on EIF3A as a direct target of MSI2 that mediates its downstream effects.

**Figure 6.**
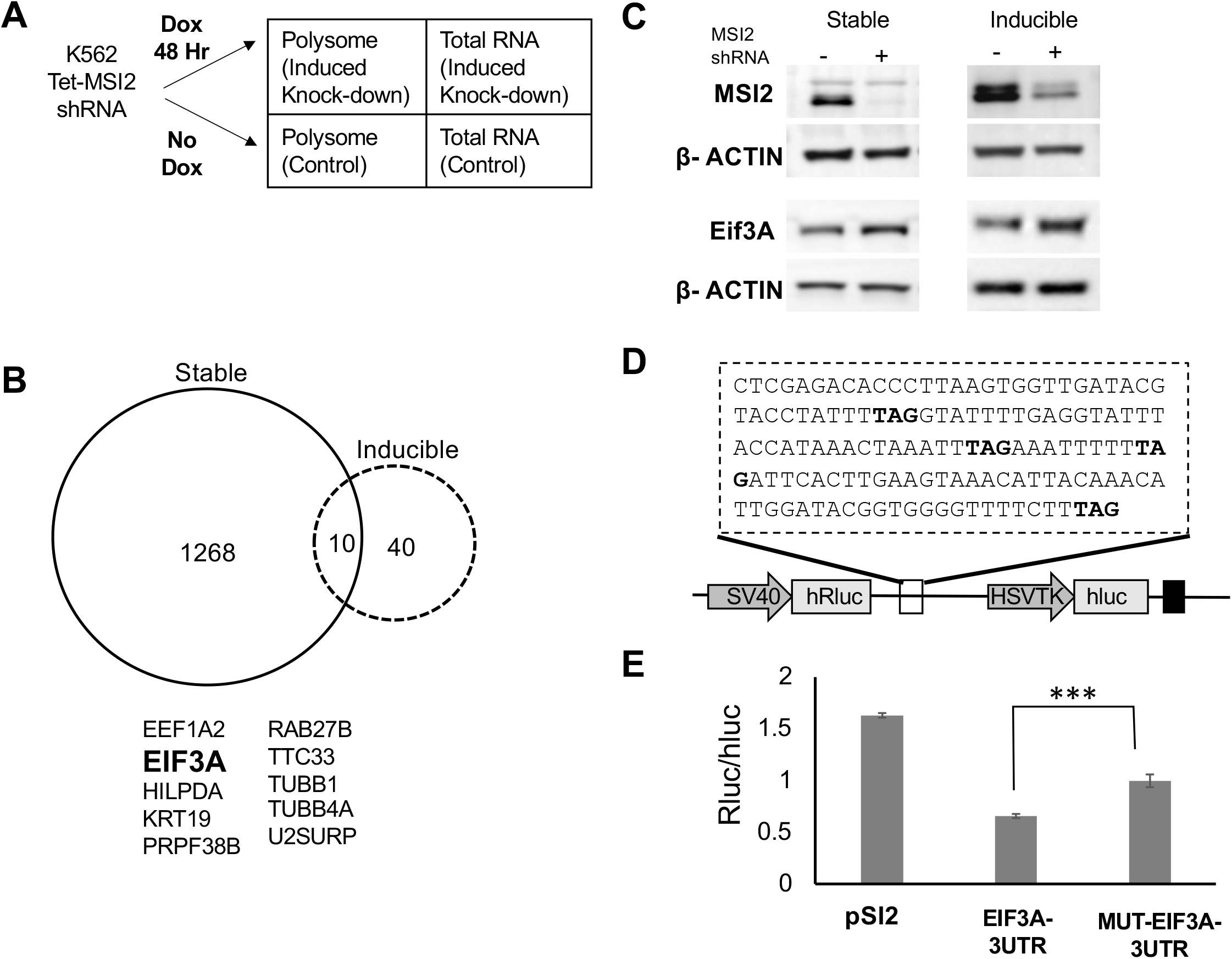
Polysome analysis and luciferase assay highlights EIF3A to be a pronounced MSI2 target. (A) Experimental scheme for polysome analysis for MSI2 inducible knockdown (similar to Figure 3B).RNA-Seq was performed on paired polysomal RNA and total RNA from induced and uninduced MSI2-KD cells after 48hrs of doxycycline induction (in two biological replicates) (B) Overlap of the genes from the polysome dataset of stable and inducible MSI2-KD cells shows 10 genes to be changing in both the stable and inducible knockdown of MSI2. EIF3A was noted to be upregulated in both conditions (2-fold in inducible and 6-fold in stable). (C) Western blot showing loss of MSI2 in the stable and inducible MSI2-KD and upregulation of EIF3A levels in both the stable and inducible MSI2-KD along with the loading control (β-actin). (D) 3’UTR region of EIF3A transcript cloned in the psicheck-2 vector downstream of Renilla luciferase (hRluc, driven by SV40 promoter). The vector also incorporates firefly luciferase (hLuc) driven by HSVTK promoter as internal control. TAG highlighted in bold was mutated to TCG to abrogate MSI2 binding. (E) Luciferase activity (normalized to fire fly luciferase) of various constructs in psicheck2 vector. These vectors were co-transfected with pCDNA-MSI2. These include pSI2 (without any insert), EIF3A 3’UTR and mutant EIF3A 3’UTR. Transfections were performed in NIH3T3 cells and luciferase activity measured 48 hours later. ** denotes p value of 0.0019. Expression of mutant 3’UTR of EIF3A resulted in increased Renilla to firefly luciferase activity as compared with the unmutated control and the empty vector backbone (control) in NIH3T3 cells.

We first confirmed the change in EIF3A expression at the protein level upon MSI2 knockdown in both inducible and stable knockdown cells (**Figure 6C**). Next, to determine if functionally repressive MSI2 binding to EIF3A 3’UTR transcript could be demonstrated, we cloned the 3’UTR region of the EIF3A transcript with iCLIP peaks into the psiCheck-2 vector downstream of the renilla luciferase construct (**Figure 6D**). Mutant 3’UTR (TCG instead of TAG, the minimal MSI2 binding motif) was generated by site-directed mutagenesis. The constructs were co-transfected with MSI2 expression vectors (pCDNA-MSI2) into NIH-3T3 cells (chosen given their lack of expression of MSI2) (24). As shown in **Figure 6E**, expression of mutant 3’UTR resulted in an increase in luciferase activity compared to the wildtype control, suggesting direct binding of MSI2 to the 3’UTR of EIF3A.

After demonstrating that EIF3A a direct target of MSI2 in K562 cells, we then sought to determine if some of the effects of MSI2 on peptide translation can be attributed to EIF3A. We hence generated inducible knockdown cells of EIF3A using the doxycycline inducible lentiviral vectors expressing two independent shRNA constructs. We first verified knockdown of EIF3A upon doxycycline induction (both for transcript and protein, **Figure 7A and Supplementary Figure 6A-C**). Accordingly, we selected 72 hours of induction as the optimum duration of doxycycline induction and performed polysome profiling of EIF3A knockdown cells. We found that similar to MSI2, knockdown of EIF3A predominantly affects the polysome (1125 transcripts**)(Figure 7B**) compared to transcripts (140 transcripts**) (Figure 7C and Supplementary File3**). Finally, to test our hypothesis that EIF3A is a critical downstream mediator of MSI2, we then determined the overlap of polysome changes in stable MSI2 knockdown to those acutely induced by EIF3A. 38 transcripts were found to overlap between the two datasets, which when analyzed using Fisher’s exact test was found to be highly significant ( p < 2.2 e-15), as shown in **Figure 7D (full list is provided in Supplementary File 4**).We also compared the changes at the protein level with acute loss of EIF3A with β-actin as the loading control (**Supplementary Figure 7**). Taken together, we conclude that EIF3A is a direct target of MSI2, and its down-regulation by MSI2 can account for a significant proportion of its indirect targets.

**Figure 7.**
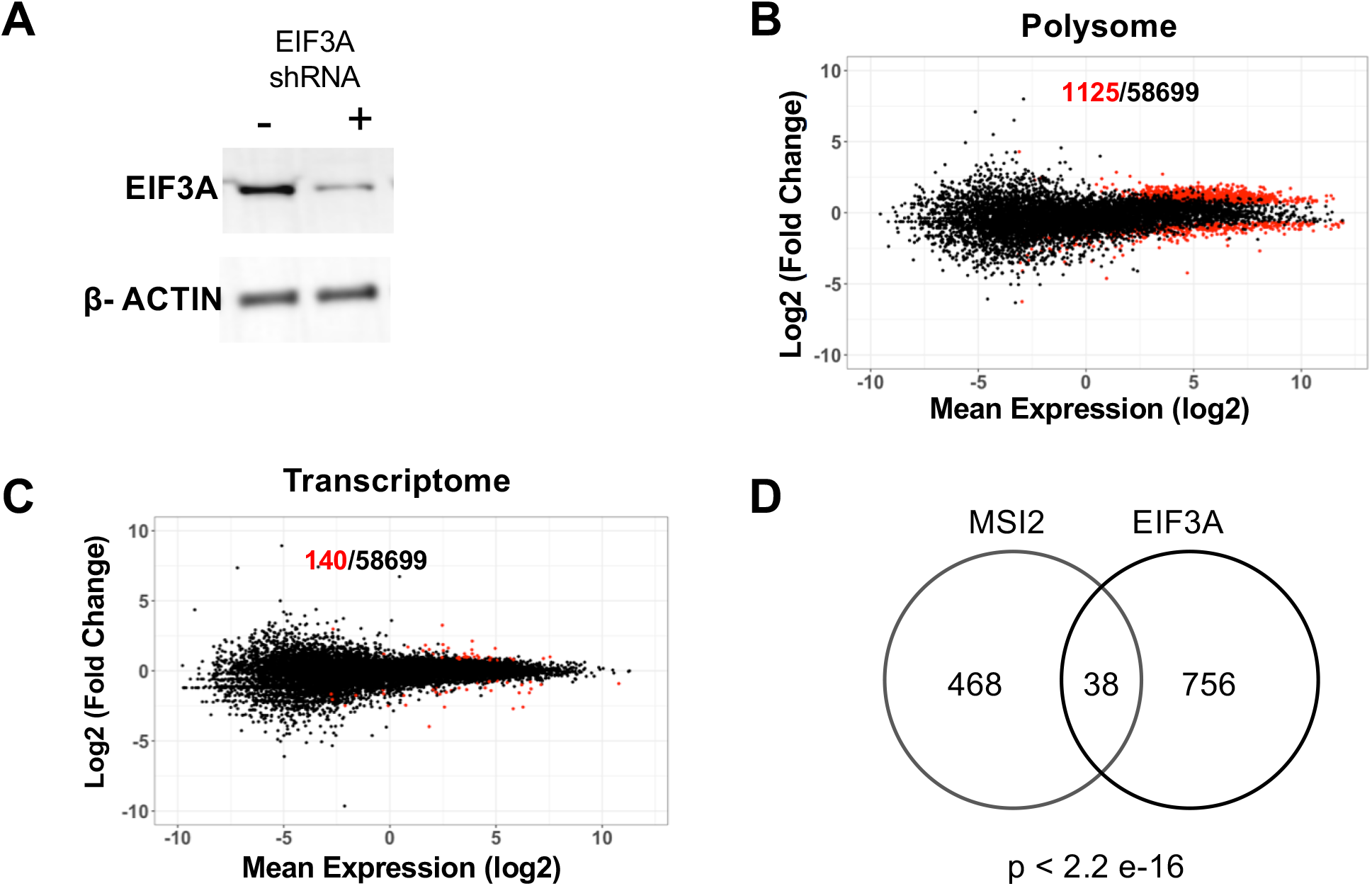
Polysome and transcriptome analysis after EIF3A inducible knockdown. (A) Western blot of uninduced and induced EIF3A knockdown cells after 72hrs of doxycycline treatment showing significant loss of EIF3A levels with anti EIF3A shRNA. (B) MA plot (Log_2_ mean expression plotted against log_2_ mean fold change) for polysome profiling of uninduced and induced EIF3A knockdown after 72hrs. Red dots indicates the transcripts that changed significantly, and the black dots represent the total number of transcripts identified. (C) MA plot (Log_2_ mean expression plotted against log_2_ mean fold change) for transcriptome from polysome profiling of uninduced and induced EIF3A knockdown after 72hrs. Red dots indicates the transcripts that changed significantly and the black dots represent the total number of transcripts identified. (**D**) Overlap between polysome profiling of stable knockdown of MSI2 and those of inducible knockdown of EIF3A. 38 overlapping transcripts were shown to overlap between the two datasets. P value calculated with Fisher’s exact test.

## DISCUSSION

In this study, we have explored the molecular mechanisms that underlie translational repression by MSI2 in myeloid leukemia cells. Up-regulation of both Musashi homologues MSI1 and MSI2 have been reported in numerous neoplasms including aggressive myeloid leukemia (43). Previous studies have used CLIP and SELEX to define sequence specificity of Musashi binding (10,14,16,30–32) and these studies generally agreed on two important findings: (1) several hundred to thousands of targets are bound by Musashi proteins (14,30) and (2) binding regions are enriched for the trinucleotide motif U/TAG (16,32). These results are in contrast with previous reports that focused on few select targets of functional significance, such as the Notch inhibitor NUMB and cyclin-dependent kinase inhibitor p21^Cip1^ (31,44). Binding of MSI1 and MSI2 to thousands of targets enriched by the U/TAG motif was surprising given their relatively narrow physiological role to regulate asymmetric cell division and quiescence. Here, we integrated two genome-wide approaches (iCLIP and polysome profiling) to test the hypothesis and identify specific functional targets of MSI2.

Our iCLIP analysis revealed over 4000 high confidence RNA binding partners of MSI2 in K562 cells with enrichment for U/TAG and poly-U/T containing pentamers, in agreement with previous results (11,13, 14). In contrast, polysome analysis of cells with stable knockdown of MSI2 showed that discernible changes at level of translation is seen only in a small fraction (2.6%) of the transcripts bound by MSI2 per iCLIP. Additionally, changes in translation were more pronounced than those in the transciptome (only 0.46%), confirming the primary role of MSI2 as a translational regulator.

To our knowledge, ours is the first study that employed polysome profiling for this purpose; two previous studies that employed the related technique ribosome-profiling both noted low coverage of ribosome footprints to be a limitation that likely underestimated translational changes (10,32). Given that our experiments were performed in stable knockdown cells, polysome dataset likely contains both direct and indirect targets. We presumed that direct targets of MSI2 are expected to have iCLIP peaks associated; accordingly, 54% of transcripts were translationally upregulated. In comparison, 26% of transcripts that were downregulated in response to MSI2 knockdown, had iCLIP peaks. Interestingly, about a quarter of all transcripts detected in the polysome fraction had significant iCLIP peaks. Thus, a mere change in translation after stable MSI2 knockdown cannot be interpreted as a direct effect of MSI2. The absence of iCLIP peaks in transcripts with discernible change in translation likely points to an effect of downstream mediators regulated by MSI2.

Through further in-depth analyses of the iCLIP and polysome datasets, we were able to define some parameters that distinguish RNA targets with productive binding from those bound non-productively. Productive targets had higher U/TAG content of crosslink clusters as well as higher total number of clusters and cluster density within the transcript coordinates. We suspect that our analysis was somewhat constrained by the presence of both direct and indirect targets within these datasets, because about a quarter of all polysome-associated transcripts had high confidence cross-link clusters while slightly more than half of the upregulated transcripts have similar iCLIP clusters. The wide variation of U/TAG content and cluster density suggests that other factors, such as additional proteins or a specific cell context, may modulate targeting. Despite this limitation, the highly significant differences between the subsets support the notion that productive binding of MSI2 likely involves multiple MSI2 binding events on each affected targets.

To definitively identify direct targets of MSI2, we performed polysome profiling after acute depletion of MSI2 (through doxycycline-inducible shRNA). Notably, far fewer transcripts (50) were noted to change on acute MSI2 loss. Of these, we chose to pursue EIF3A given it was one of the few transcripts with sustained upregulation upon stable MSI2 knockdown. In addition to its role in canonical initiation of translation, EIF3 is also now known to regulate specific transcripts that regulate cell proliferation (42). After confirming a change to EIF3A protein expression and verifying direct binding of MSI2 to 3’UTR (luciferase assay), we performed polysome profiling of EIF3A knockdown cells and intersected the results with MSI2 stable knockdown. These results showed significant overlap (p-value < 2.2e-16) between EIF3A targets and MSI2 targets. Our results strongly suggest a role for EIF3A in mediating the broad effects of MSI2 on the polysome. Importantly, EIF3A expression is dysregulated in multiple cancers (both up and down regulated) (41,45–48). Taken together, our results suggest that biological function of MSI2 may be mediated in part through another critical translational regulator, EIF3A. While the overlap of EIF3A and MSI2 targets were highly significant, majority of transcripts were not shared between the datasets. This suggests that at steady state, downstream effects of MSI2 are likely complex and not attributable to a single mediator. EIF3A is also a critical component of the canonical EIF3 complex and its drastic down-regulation (as achieved by shRNA) likely leads to other physiological effects unrelated to MSI2 mediated suppression. It also needs to be determined if the MSI2-EIF3A connection is cell-context specific, and operant in MSI2-mediated oncogenesis.

Finally, our results also highlight overall challenges in determining functional targets of RBP. Cross-link immunoprecipitation has helped define the RNA targets bound by dozens of RBP, but mere binding is not synonymous with a discernible functional change. Computational algorithms that rely on intensities and positions of binding peaks have been proposed to distinguish functional binding (49). Our own results also show that RNA targets functional binding of MSI2 correlates with higher density binding to targets. Such a correlation is however imperfect and not able to distinguish such targets precisely. For MSI2, integrating a functional end point (peptide translation) distinguish these functional targets which are only a fraction of all bound targets.

## DATA AVAILABILITY AND ACCESSION NUMBERS

iCLIP analysis files (20160613_Manoj library) are available at http://icount.biolab.si. Sequencing files have been deposited in GEO (GSE93210), and reviewers may access these at the URL https://www.ncbi.nlm.nih.gov/geo/query/acc.cgi?token=avmtiwgazzolxgv&acc=GSE93210.

## SUPPLEMENTARY DATA

Supplementary data files are available at NAR online.

## ACKNOWLEDGEMENTS

We thank Jeffrey Kieft (University of Colorado Denver) and Haifan Lin (Yale) for help with polysome fractionation and helpful suggestions. The Yale Stem Cell Center’s Genomics Core provided assistance with sequencing.

## FUNDING

The work was funded in part by National Institutes of Health R01 HL104070 (to MMP), R01 HL133406 (to MMP and KMN) and a pilot grant from the Yale Cancer Center (to MMP). Core facilities used in this study were supported in part by U54 DK106857

## CONFLICTS OF INTEREST

The authors have no conflicts of interest to state.

## AUTHOR CONTRIBUTIONS

SK, OR, VB, AKG, AKM, AEM, NN and RJR performed experiments, KVP, IRdlM and MMP performed informatics analysis, MMP and KMN provided experimental guidance, MMP designed the study, SK, KMN and MMP, wrote the paper with input from other co-authors.

## Notes

https://www.ncbi.nlm.nih.gov/geo/query/acc.cgi?token=avmtiwgazzolxgv&acc=GSE93210.

http://icount.biolab.si

### Competing Interest Statement

The authors have declared no competing interest.

