## Supplementary Information for "Integrative genome-wide analysis reveals EIF3A as a key downstream regulator of translational repressor protein Musashi 2 (MSI2)"

### SUPPLEMENTARY FIGURE LEGENDS:

**Supplementary Figure 1. Optimization of iCLIP.**
A: FLAG-MSI2 expression in K562-FLAG-MSI2 cells and control cells by Western blot (anti-FLAG antibody).
B: Autoradiographs of CLIP-Seq to show association of RNA to FLAG-MSI2. Ligation of P-32 labeled 3' linker performed on K562-FLAG-MSI2 lysates digested with RNAse-A either crosslinked with UV (+) or non-cross-linked controls (-) and resolved on PAGE gels and transferred to nitrocellulose. Membranes visualized by autoradiography shows bands (including one corresponding to MSI2) in the cross-linked lane alone.
C: Optimization of RNAse conditions for iCLIP. Varying dilutions of RNAse A (of a 10U/ml stock concentration) for RNAse digestion after cell lysis per the iCLIP protocol. A dilution of 1:50 was found to give the optimal smear pattern for library preparation
D: Optimization of PCR cycles for cDNA library preparation. As per original iCLIP protocol, three regions of RNA-MSi2 was cut out per molecular weight (L: low, M: medium and H: high) to ensure adequate representation of fragments of differing size. After RT, PCR performed at 20 and 22 cycles to determine optimal cycle number. 20 cycles were used in this case given adequate product visualization at 20 cycles.

**Supplementary Figure 2.**
MSI2 expression in the stable MSI2 knockdown clones by qPCR. Results are normalized to beta-actin expression.

**Supplementary Figure 3. Probability of secondary structural conformations of sequence stretches around cross-link clusters in the 3'UTR of three groups of transcripts (unchanged, upregulated and downregulated) at the level of translation upon MSI2 knockdown.**
40 bp region (20 bp upstream or downstream of the cross-link site) was selected for high confidence cross-link sites (FDR <0.05) and sequence extracted as a FASTA file. A random set of 1000 sequences analyzed for each group. The CapR algorithm predicts probability of each base-pair to be part of 6 secondary structure conformations (Bulge, Exterior, Hairpin, Internal, Multibranch or Stem, listed in panel headings A-F). Plotted are the average probabilities for each position (cross-link site is at position 20). The distribution of probabilities was not different for the three groups plotted for any of the conformations.

**Supplementary Figure 4. Protein expression of additional targets as predicted by polysome profiling (**SP1, C-MYC, RAD21, USP28 and RB1 ) in cells with stable MSI knockdown. Beta-actin loading control is included.

**Supplementary Figure 5**. **Optimization of time point for induction by doxycycline on MSI2 inducible clones for polysome profiling.**
(A) Confirmation of knockdown of MSI2 in uninduced and induced cells on doxycycline treatment by western blot.
(B) Timeline of MSI2 knockdown with inducible shRNA for inhibiting MSI2 expression by qPCR.
(C) MSI2 depletion at 48hrs with Doxycycline from inducible clones in biological replicates determined by qPCR
(D) Representative polysome profiles of control and Doxycycline treated inducible clone of MSI2.

**Supplementary Figure 6 Optimization of time point for induction for shRNA mediated depletion of EIF3A on single cell clones.**
(A) Two different shRNA used for generating EIF3A knockdown single cell clones.
(B) and (C) Maximum effect of doxycycline induction on shRNA mediated EIF3A depletion is observed after 72 hours at both the transcript and protein levels

**Supplementary Figure 7. Altered expression of proteins in Doxycycline induced EIF3A knockdown clones determined by Western blotting.** With acute depletion of EIF3A for 72hrs CDK6, SP1, C-MYC, RAD21 were observed to be downregulated although USP28 did not show any observable change.

**SUPPLEMENTARY FILES:**

**Supplementary File 1:** Analysis of Polysome and Total RNA from MSI2 knockdown (both stable and inducible knockdown) using Cufflinks. Shown are the gene name FPKM, log2 fold change, test status, p value, q value and inferred significance of the comparison.

**Supplementary File 2: Integration of iCLIP and polysome datasets.** Gene names, overlap of iCLIP and Polysome dataset (in number of replicates) and regulation status (Upregulated, Downregulated or Unchanged) are listed in the columns

**Supplementary File 3. Polysome analysis of EIF3A inducible knockdown.**
Doxycycline Inducible knockdown of EIF3A at 72 hrs were compared with the uninduced control for Polysome associated RNA (EIF3A_KD_IND_POLY and MSI2_KD_UNIND_POLY) and the Total cellular RNA (EIF3A_KD_IND_TOTAL and EIF3A_KD_UNIND_TOTAL) using cufflinks. Each set had two biological replicates. The FPKM, log2 fold change, significance, test value, p-value and q-value of the comparison between the datasets are shown here.

**Supplementary File 4. Overlap of polysome from MSI2 stable knockdown and EIF3A inducible knockdown.** The datasets of MSI2 stable knockdown and EIF3A inducible knockdown are compared using Fishers Test to determine the common partners that are being regulated by both the translational regulators.

**SUPPLEMENTARY METHODS:**

**1. OLIGONUCLEOTIDES**

FLAG-MSI2-F
cctggatCCACCATGGACTACAAAGATGACGACGACAAGATGGCCGATCTGACATCG

FLAG-MSI1-R cctgtcgaCGGGACTGTGTCCTCTCTCTC

MSI2 QPCR-F- GTTATCTGCGAACACAGTAGTG

MSI2 QPCR-R- ACCCTCTGTGCCTGTTGGTAG

EIF3A-PSICHECK2-F- tcgctcgagACACCCTTAAGTGGTTGATACG

EIF3A-PSICHECK2-R- agagcggccgcAGCAGCAAGTTCTGTGTTGG

EIF3A Q5SDM-PSICHECK2-F – CTGGCCGCAATAAAATATC

EIF3A Q5SDM-PSICHCEK2-R- TACTGTTTGGATTCACAC

QRT-EIF3A-F- AGCCTGCCGCCAAGATG

QRT-EIF3A- R- GCAGGCTGCTTTTTGCCA

**2. shRNA CLONES:**

| Clone ID | Sequence of insert (5'→3') |
|---|---|
| MSI2-9 | CCGGGTGGAAGATGTAAAGCAATATCTCGAGATATTGCTTTACATCTTCCACTTTTTG |
| MSI2-11 | CCGGCCCAACTTCGTGGCGACCTATCTCGAGATAGGTCGCCACGAAGTTGGGTTTTTG |
| MSI2-12 | CCGGAGATAGCCTTAGAGACTATTTCTCGAGAAATAGTCTCTAAGGCTATCTTTTTTG |
| Control | CCGGCAACAAGATGAAGAGCACCAACTCGAGTTGGTGCTCTTCATCTTGTTGTTTTTG |
| EIF3A-7 | CCGGGCGCCTTGAGAGTCTGAATATCTCGAGATATTCAGACTCTCAAGGCGCTTTTTG |
| EIF3A-27 | CCGGCGTGCTGATGATGATCGGTTTCTCGAGAAACCGATCATCATCAGCACGTTTTTG |

**3. POLYSOME PROFILING (**Adapted from Merrick et al (1))

Sucrose gradient
10-60% sucrose gradient was prepared in 5 ml centrifuge tubes from the base solutions (recipes below) the previous day using the Teledyne ISCO-160 Gradient Former (Lincoln, NE) and allowed to further linearize overnight at 4ºC.

Cell culture and harvest
40-50 million cells in their log-phase of growth were used per sample (growing at density of 0.5-0.75 million cells/ml in two T75 ml flasks). Media changed daily for three days leading to the day of analysis to ensure log-phase growth.

On day of experiment, Cycloheximide (Sigma, C0934) was added to the cultures at final concentration of 100 ug/ml to the cells and returned to 37°C incubator for 10 minutes to induce translational arrest. Cells were harvested rapidly at end of the 10 minute incubation by spinning at 1000 rpm for 5 minutes at 4°C and washing once with ice-cold PBS + cycloheximide (100ug/ml). Total RNA from 1-2 million cells was isolated for mRNA-Seq. Cells were lysed with with 700-900 ul of TMK100 buffer and lysates cleared by spinning at max speed (~ 13000 rpm) on standard tabletop micro centrifuge at 4ºC. Cleared lysates were loaded to the top of the centrifuge tubes and subjected to ultracentrifugation (3 hours at 35,000 RPM on a Beckman SW40 Ti rotor). Fractions were collected immediately after centrifugation using the Teldyne ISCO automated fraction collector with continuous monitoring of absorbance at 254 nm. A total of 20-46 fractions were typically collected per sample (800 ul each) depending on the elution profile.

<u>RNA-isolation, library preparation and sequencing</u>
100 ul of fractions corresponding to heavy polysomes per absorbance tracing (#26-39) for the MSI12 stable-knockdown cell line, (#12-17) of inducible MSI2 knock-down and (#11-16) for inducible EIF3A knockdown line was pooled and diluted 1:1 with RNA-seq free water. Total polysomal RNA was isolated using Trizol (Life Technologies, Carlsbad, CA). Polyadenylated RNA was isolated using magnetic oligo dT beads (New England Biolabs, Ipswich, MA) and multiplexed mRNA libraries prepared using Illumina Truseq RNA-Seq kits. Libraries were similarly prepared from total RNA isolated at the same time. Sequencing was performed on Illumina HiSeq2000 platform (50 base pair single end).

**Buffers:**

<u>TMK-100 (Tris/Mg2+/K+) lysis buffer</u>
10 mM Tris-Cl, pH 7.4
5 mM $MgCl_2$
100 mM KCl
1% (v/v) Triton X-100
0.5% (w/v) deoxycholate
Prepare in nuclease free (DEPC or molecular grade) water.

Right before use, add:
1 U/ml ribonuclease inhibitor (like RNAsin from Promega)
2 mM dithiothreitol

<u>Sucrose gradient solutions, 10% and 60% (w/v)</u>
10% or 60% (w/v) sucrose
100 mM KCl
5 mM $MgCl_2$
2 mM dithiothreitol
20 mM HEPES-KOH, pH 7.4
Prepare in nuclease free water (DEPC or molecular grade). Add 1 U/ml ribonuclease inhibitor (like RNAsin from Promega) and 2 mM dithiothreitol right before use.

#### 4. EIF3A sequence

CTCGAGACACCCTTAAGTGGTTGATACGTACCTATTT**TAG**GTATTTTGAGGTATTTACCATAAACTAAAT
TTAGAAATTTTT**TAG**ATTCACTTGAAGTAAACATTACAAACATTGGATACGGTGGGTTTTCTT**TAG**ATT
TTACTTGAGAGAAGGTGAGTACAAAGCAATTTGCAGTTGTTGTAATGACAAGATTACTGCGCAAGTGTGA
ATCCAAACAGTA**TAG**CTTTTAAATTTTAAAGCATTTGGTAAATTATCGCTGAGTTTTTTTCTGTTGCCAA
**TAG**CAAACTGCTTTTCCATTAATGGAGAATTCATGCCTTTCAAGCATTTTAAATATGACAATATTTATAA
ATGTATGGTTTGGAGGAATCGTTTAAATTCTCTTTCCTAATTTTCTTTCTTTTGAAGA**TAG**ATTCTTTCA
ACAAGTAATTTG**TAG**TAATGACTGTGTTGACTTCAATTTTGGAGCGCAG**TAG**CTATGTTAAAGATGAACT
ATTTGGTCTCATTGAAGCCAACACAGAACTTGCTGCTGCGGCCGC

**EIF3A mutant sequence**

CTCGAGACACCCTTAAGTGGTTGATACGTACCTATTT**TCG**GTATTTTGAGGTATTTACCATAAACTAAAT
TTAGAAATTTTT**TCG**ATTCACTTGAAGTAAACATTACAAACATTGGATACGGTGGGTTTTCTT**TCG**
ATTTTACTTGAGAGAAGGTGAGTACAAAGCAATTTGCAGTTGTTGTAATGACAAGATTACTGCGCAAGTG
TGAATCCAAACAGTA**TAG**CTTTTAAATTTTAAAGCATTTGGTAAATTATCGCTGAGTTTTTTTCTGTTGC
CAA**TAG**CAAACTGCTTTTCCATTAATGGAGAATTCATGCCTTTCAAGCATTTTAAATATGACAATATTTA
TAAATGTATGGTTTGGAGGAATCGTTTAAATTCTCTTTCCTAATTTTCTTTCTTTTGAAGA**TAG**ATTCTT
TCAACAAGTAATTTG**TAG**TAATGACTGTGTTGACTTCAATTTTGGAGCGCAG**TAG**CTATGTTAAAGATGA
ACTATTTGGTCTCATTGAAGCCAACACAGAACTTGCTGCTGCGGCCGC

Q5SDM_F- CTGGCCGCAATAAAATATC
Q5SDM_R TACTGTTTGGATTCACAC

**5.WESTERN BLOT:**

| Primary Antibody | Vendor | Cat # | Host | Concentration | Secondary |
|---|---|---|---|---|---|
| C-MYC | CST | 9402 | R | 1-1000 | GαR |
| MSI2 | Millipore | 04-1069 | R | 1-1000 | GαR |
| RAD21 | SC | sc-271601 | R | 1-1000 | GαR |
| SP1 | SC | sc-59 | R | 1-1000 | GαR |
| USP28 | SC | sc-79312 | R | 1-1000 | GαR |
| CDK6 | CST | 13331 | R | 1-1000 | GαR |
| EIF3A | CST | 2538 | R | 1-1000 | GαR |
| BETA-ACTIN | CST | 4970 | R | 1-1000 | GαR |

**Abbreviations:**
**CST** **Cell Signaling Technologies**
**SC** **Santa Cruz**
**R** **Rabbit**
**M** **Mouse**
**G** **Goat**
**S** **Sheep**

All primary antibody incubations were performed overnight at 4°C. All secondary antibody incubations were performed at room temperature for 1 hour.

**6. BIOINFORMATIC ANALYSIS:**

**A. Translatome data analysis**
The three replicates of RNA-Seq data (single-end 50 bp) obtained for control and MSI2-KD samples from total and polysome fractions were preprocessed for adapter removal and quality control with Trimmomatic (2) and FastQC (http://www.bioinformatics.babraham.ac.uk/projects/fastqc/) respectively, and mapped to human

genome (UCSC h19) using STAR aligner (3). Differential expression of annotated genes (UCSC hg19) between MSI2-knockdown and Control in total and polysome fraction was calculated using Cuffdiff (4) with following additional criteria: (1) For high-confidence, minimum FPKM of genes was set to 5 for control and/or MSI2-knockdown samples. (2) Translationally upregulated genes had $\log_2$FC $\geq$ 0.58 (or 1.5 FC) in polysome fraction and $\log_2$FC $\leq$ 0 in total fraction and the ratio of $\log_2$FC for that genes between polysome and total had 2-fold difference (ie $\log_2$FC polysome – $\log_2$FC total > 1).(3) For translationally downregulated genes, expression levels in opposite direction was applied. Translationally downregulated genes had $\log_2$ FC $\leq$ -0.58 (or - 1.5 fold) in polysome and the ratio of $\log_2$FC for that genes between polysome and total had 2-fold difference (i.e. $\log_2$FC polysome – $\log_2$FC total < -1). Genes showing upregulation or downregulation on translation level in MSI2-Knockdown cells were analyzed for the enrichment of specific disease and functions by Ingenuity Pathway Analysis (IPA) (5).

**B. iCLIP data analysis**

The three replicates of iCLIP-Seq data were analyzed using iCount online server (http://icount.fri.uni-lj.si/) to determine cross-link sites and cross-link clusters of high confidence (FDR <0.05) as well as pentamers and their z-scores (6). For reproducibility, the crosslink clusters (or peaks) that had more than 50% overlap in at least two replicates were determined by bedtools (7) “intersect” function. These reproducible clusters were mapped to reference genes (protein-coding, non-coding and intergene) and transcript features (5’/3’ UTR, CDS, exon and intron) available from UCSC version hg19 annotation using bed tools. Additional motif analysis was performed using HOMER (8) for enrichment of enrichment of motifs 5-10 bp long.

As an alternate approach to determine enrichment of MSI2 binding, read density was calculated as Read Per Kilobase (RPKB) in specific regions of transcripts from mapped read files (BAM). Before mapping, the duplicate reads with 9 nts long UMI (Unique Molecular Identifier) in fastq files were collapsed using FASTX-Toolkit (http://hannonlab.cshl.edu/fastx_toolkit/index.html), and rRNA-mapping reads were discarded with Bowtie2 (9) using rRNA reference build. The remaining reads were mapped to hg19 genome using STAR version 2.4.2a. For most RPKB calculations, the BAM files from three replicates were merged, as the data was reproducible across replicates. Custom Perl scripts were used for other downstream analysis (including distribution of UAG and poly-T motifs, distribution of total cluster number, cluster density, cross-link density and TAG density). These scripts have been uploaded to Github (https://github.com/pillailab).

Analysis of secondary structures of regions around cross-link sites was performed using the CapR algorithm. A region 25 bp upstream and downstream of high confidence cross-link sites (FDR<0.05) was extracted using MEME suit (10). Relative probability of each of the 50 nucleotides to form secondary structures (Bulge, Hairpin, Multibranch, Exterior, Internal or Stem) was computed using the CapR algorithm (11). Values were averaged and plotted for each position **(Supplementary Figure 3)**.

**Supplementary Figure 1**

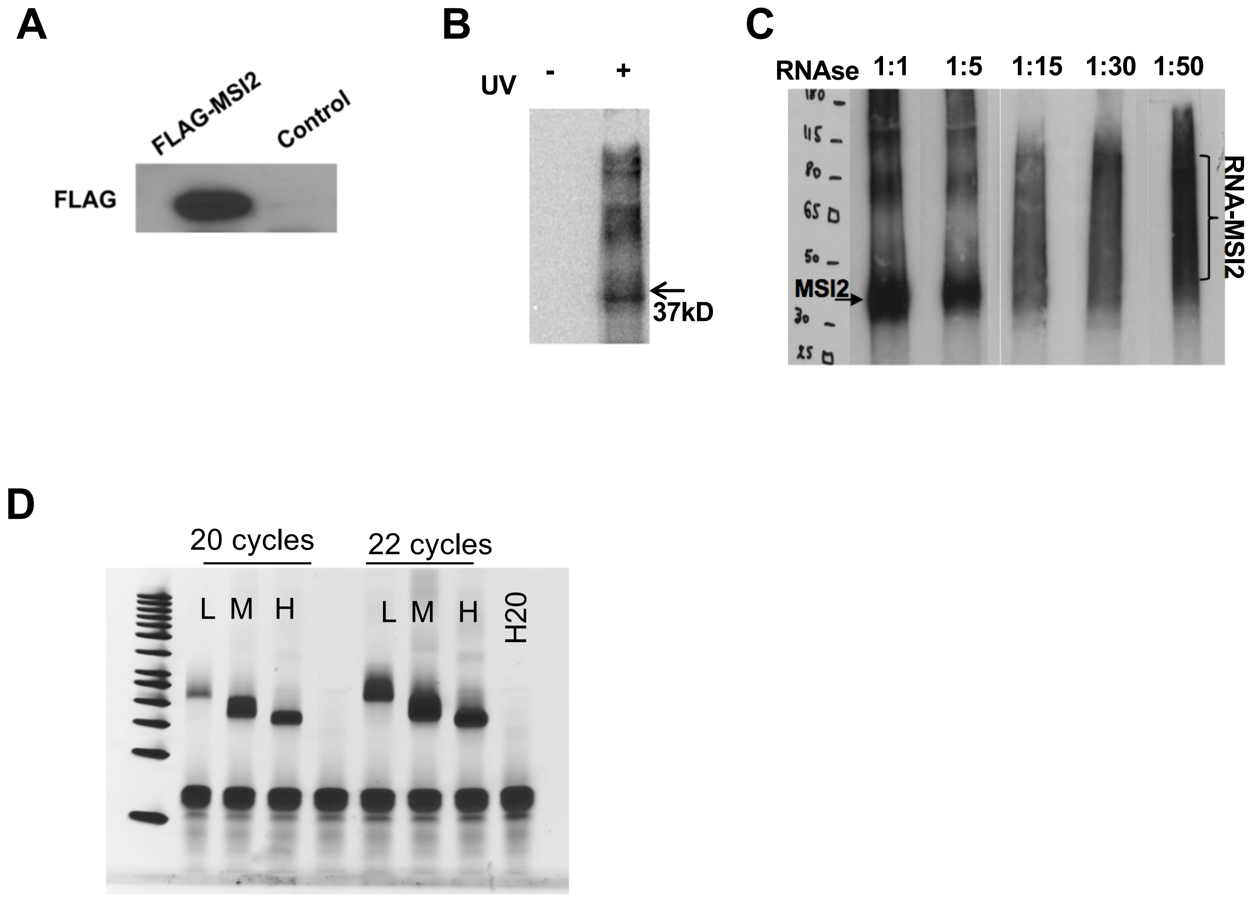


**iCLIP PCR**

**Supplementary Figure 2**

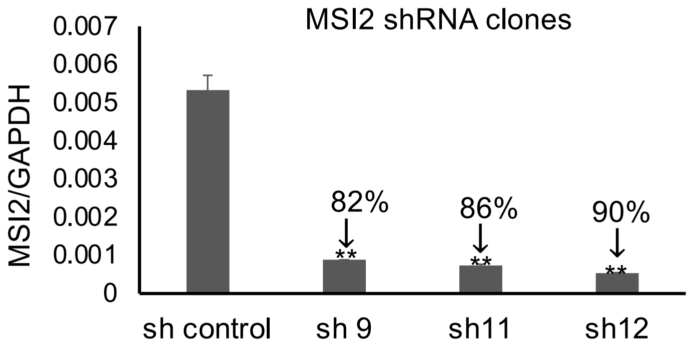

**Supplementary Figure 3**

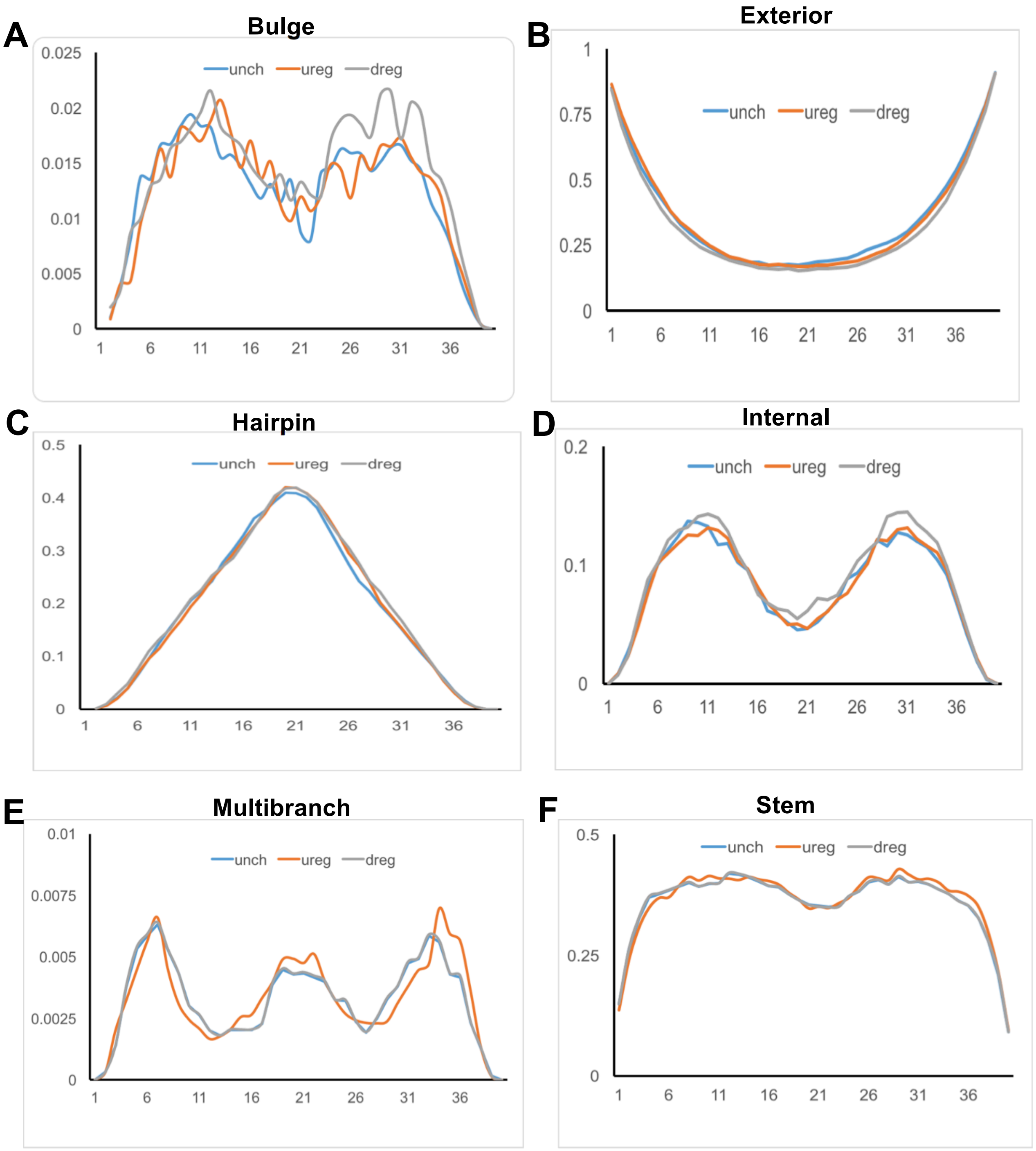

**Supplementary Figure 4**

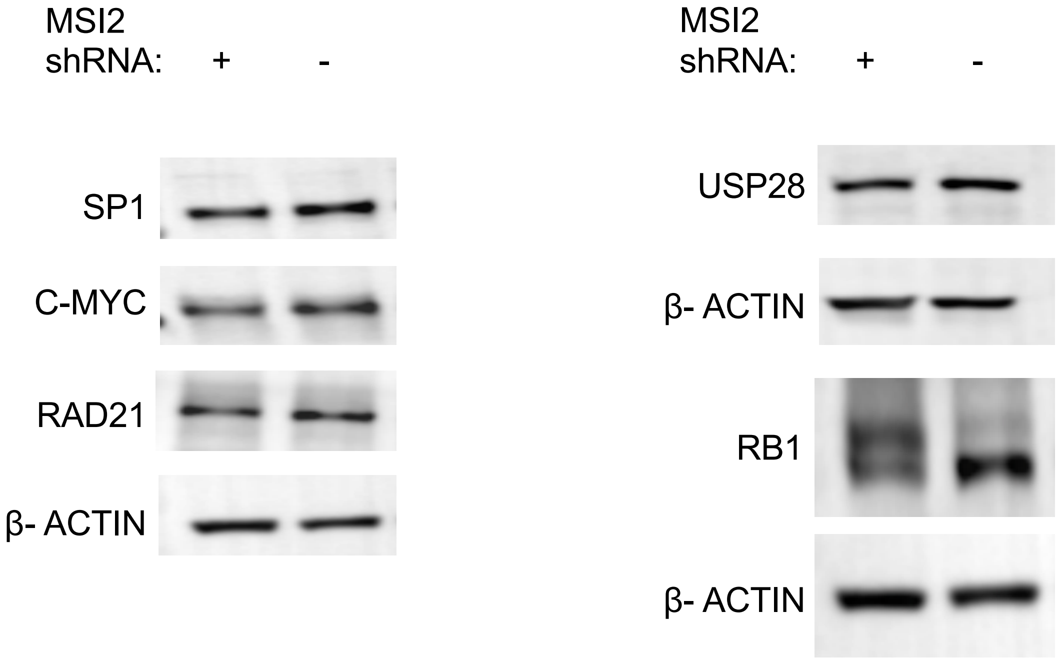

**Supplementary Figure 5**

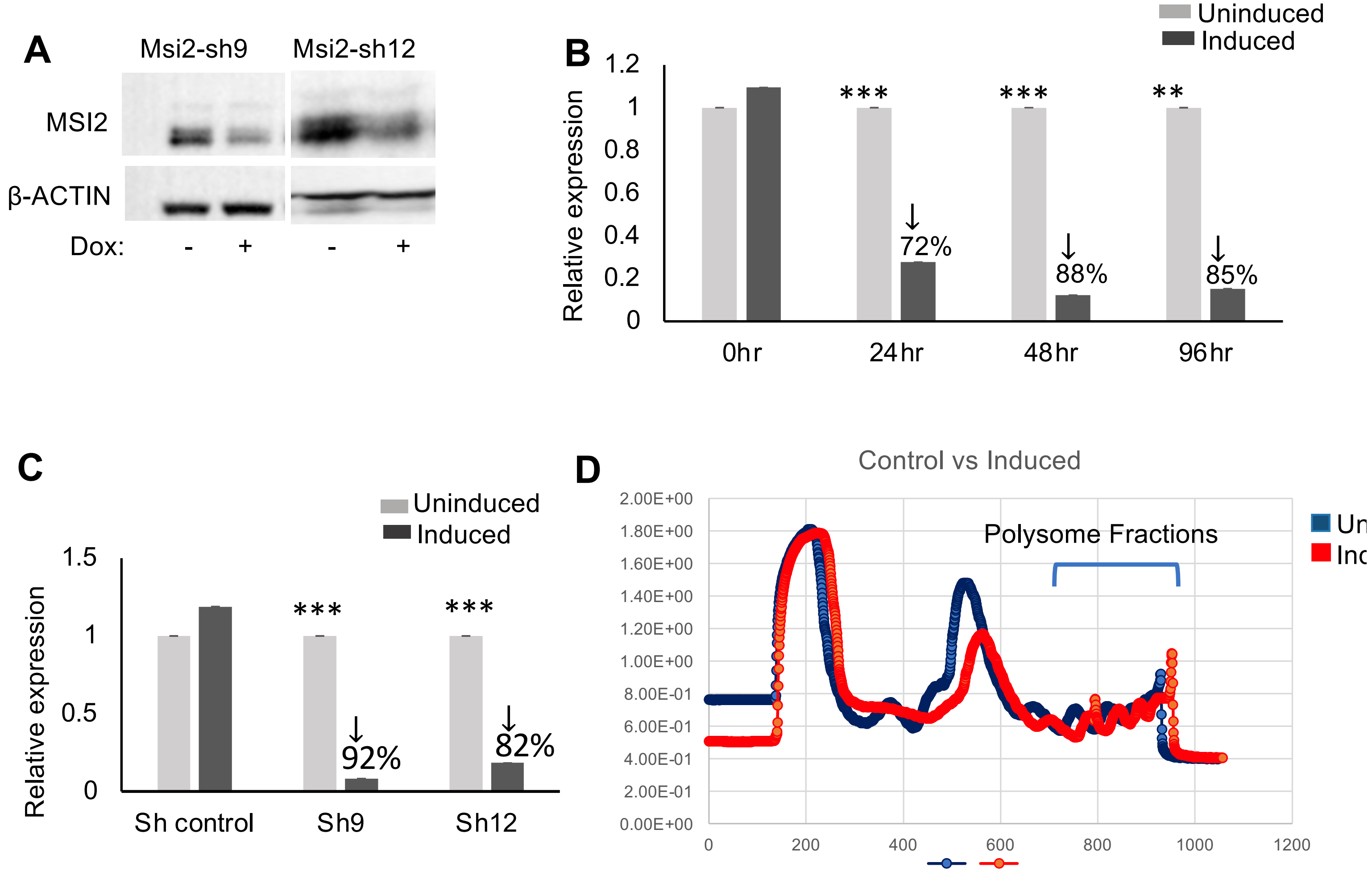

**Supplementary Figure 6**

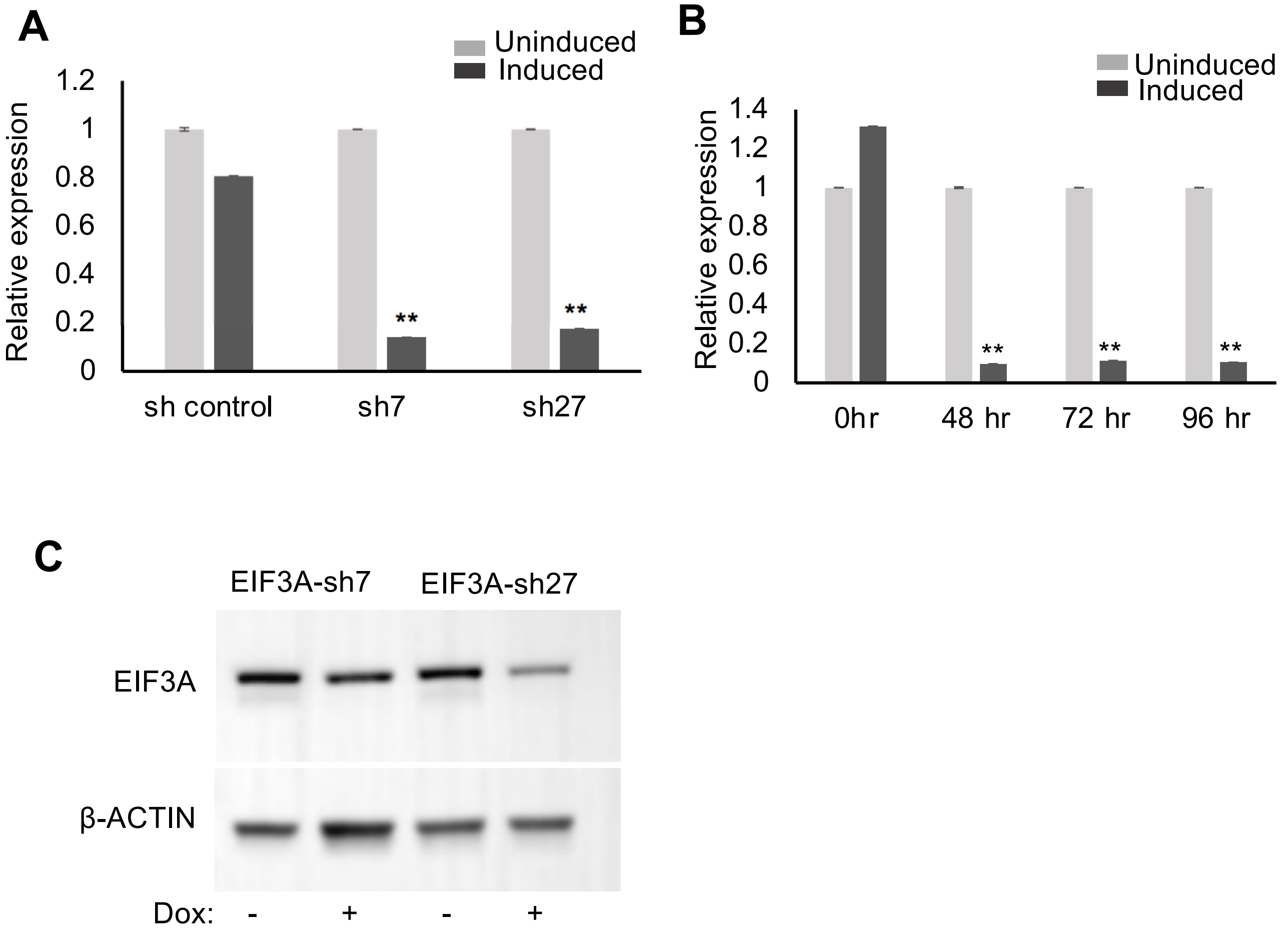

**Supplemental Figure 7**

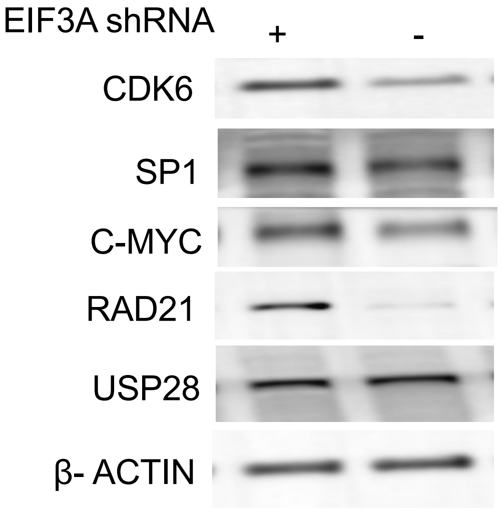
