## Supplementary Figures for "Integrative genome-wide analysis reveals EIF3A as a key downstream regulator of translational repressor protein Musashi 2 (MSI2)"

**Supplementary Figure 1**

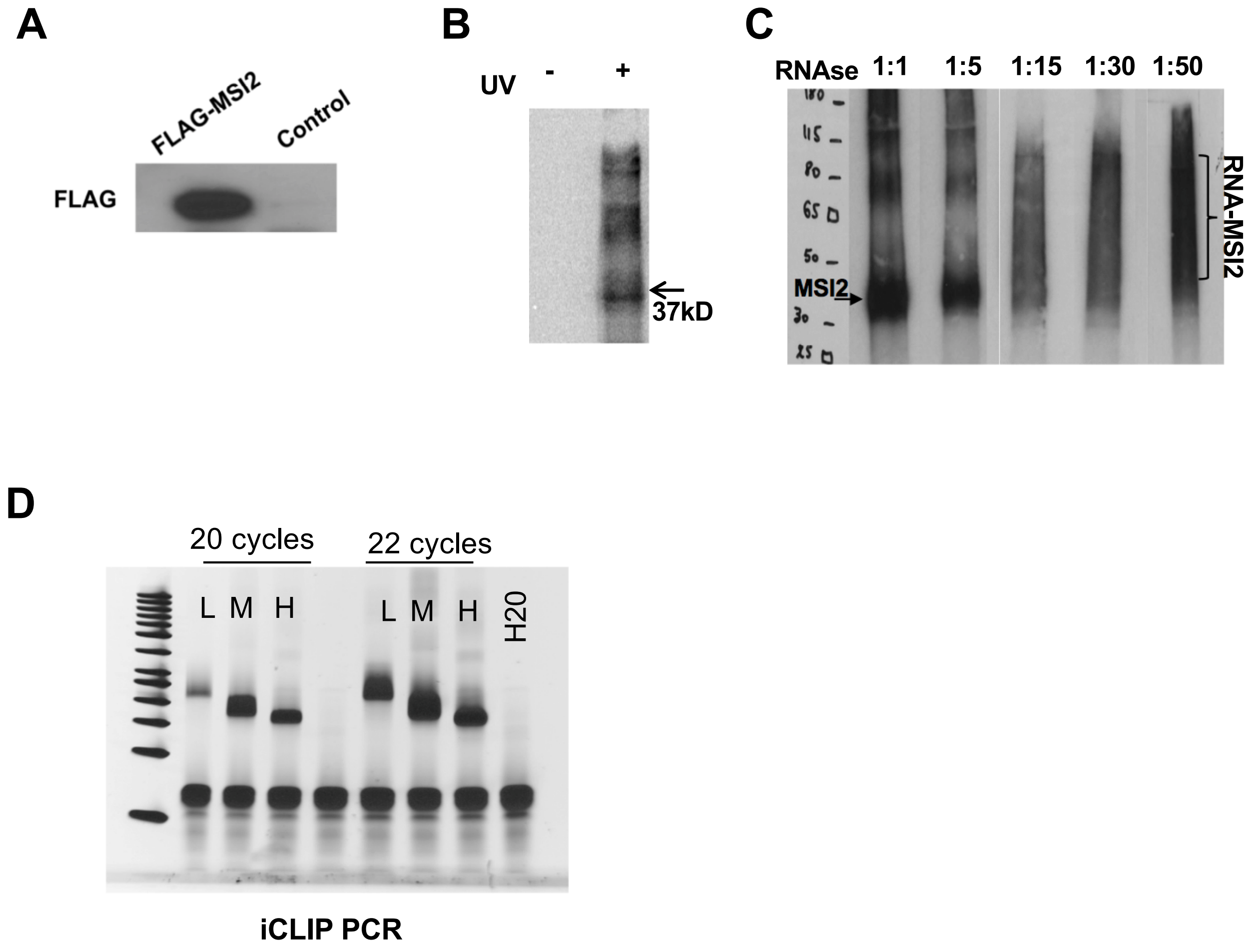

**Supplementary Figure 2**

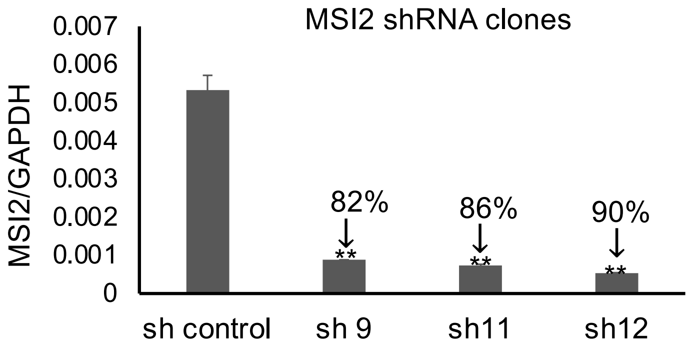

**Supplementary Figure 3**

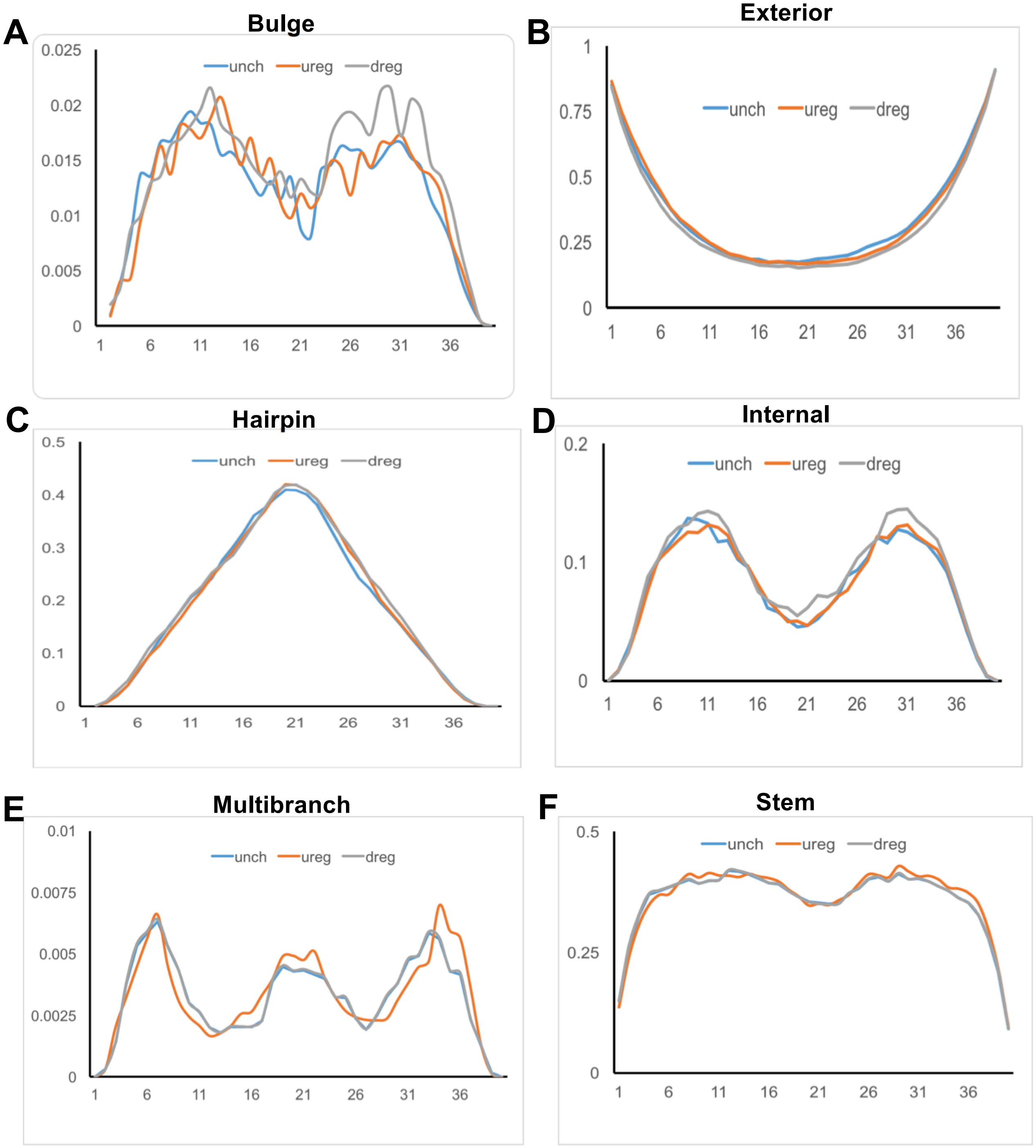

**Supplementary Figure 4**

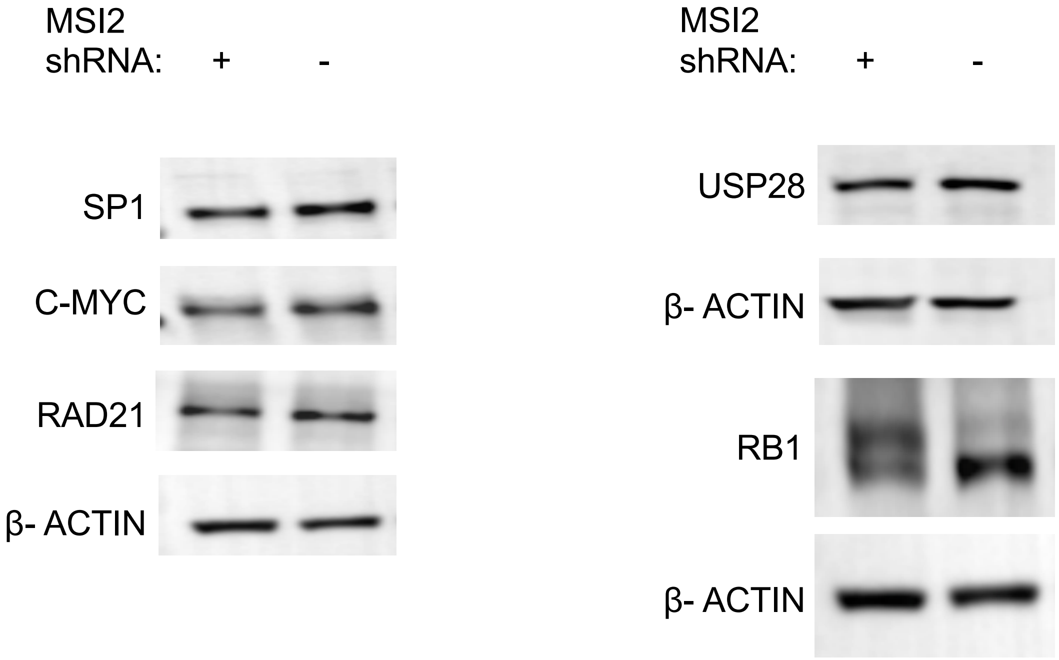

**Supplementary Figure 5**

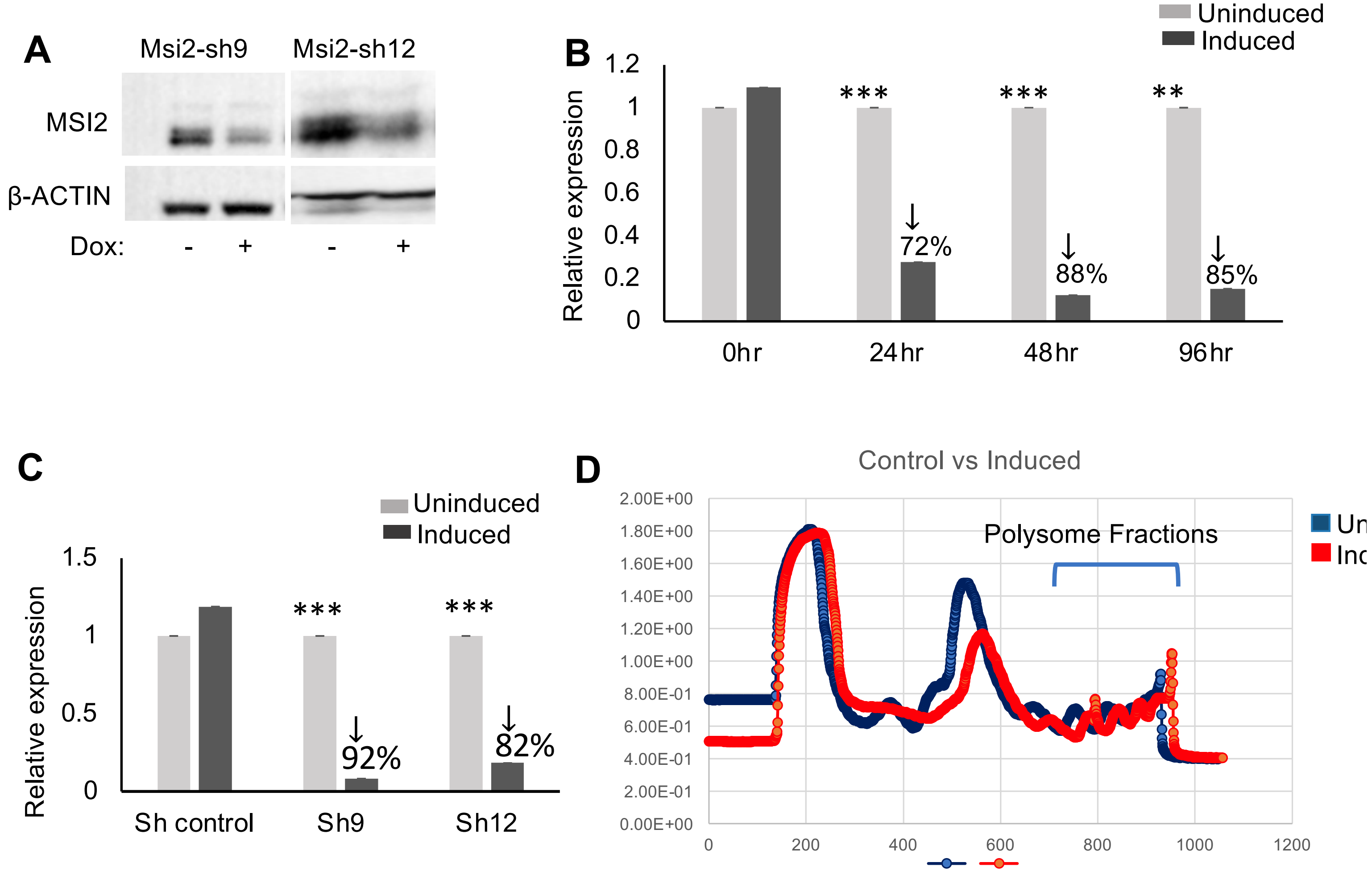

**Supplementary Figure 6**

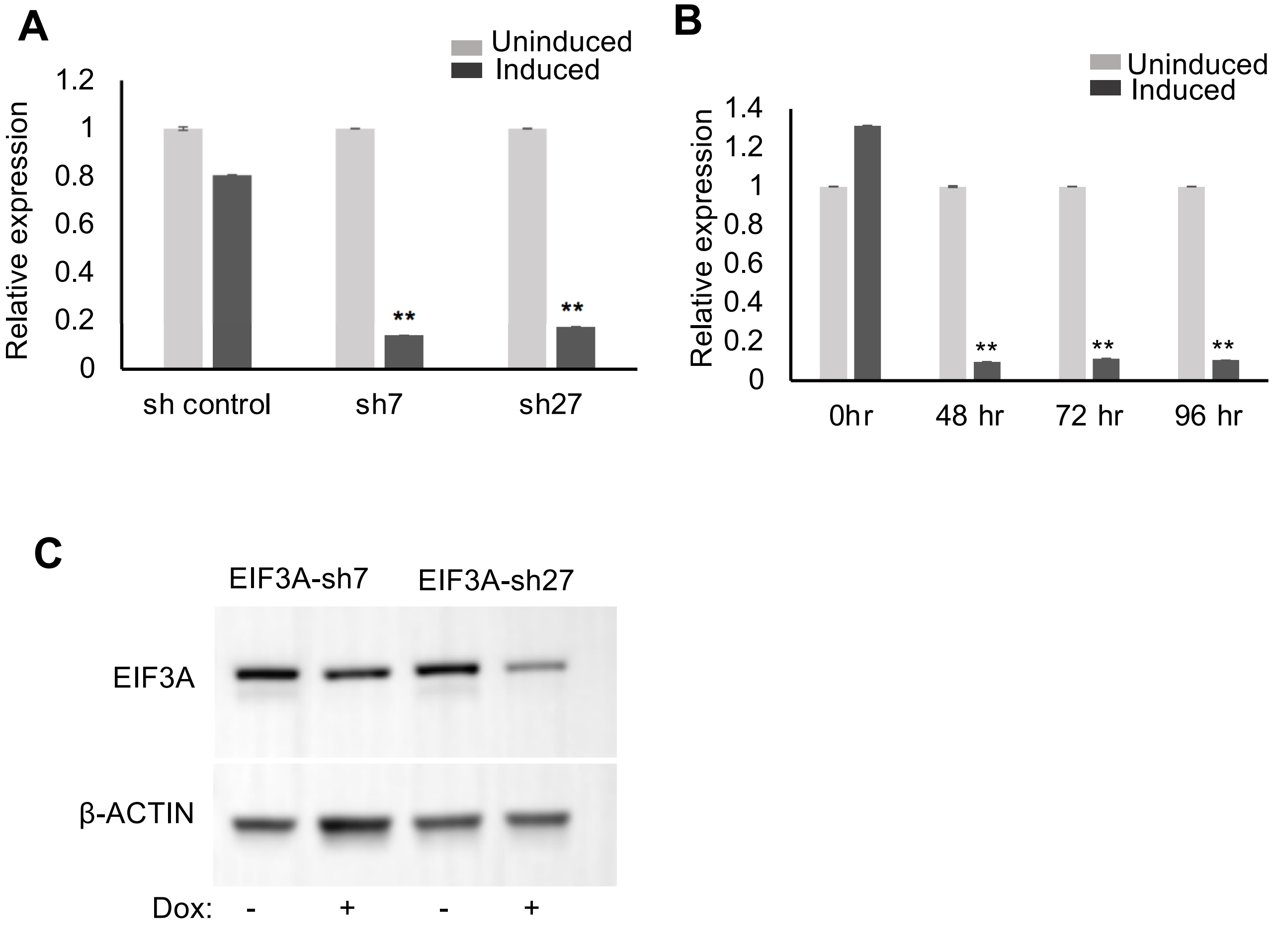

**Supplemental Figure 7**

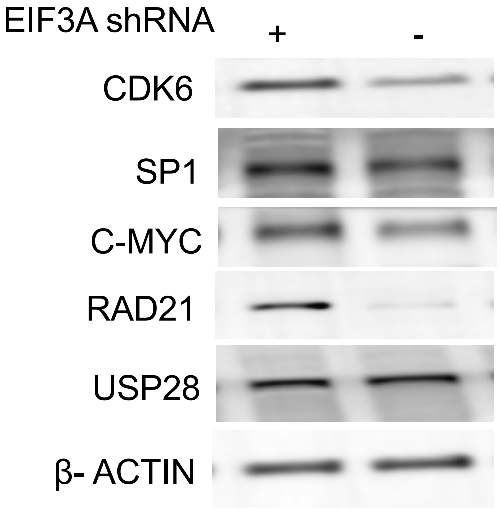
